# Sleep spindle maturation enhances slow oscillation-spindle coupling

**DOI:** 10.1101/2022.09.05.506664

**Authors:** Ann-Kathrin Joechner, Michael A. Hahn, Georg Gruber, Kerstin Hoedlmoser, Markus Werkle-Bergner

## Abstract

The synchronization of canonical fast sleep spindle activity (12.5-16 Hz) precisely during the slow oscillation up peak is considered an essential feature of adult non-rapid eye movement sleep. However, there is little knowledge on how this well-known coalescence between slow oscillations and sleep spindles develops. Leveraging individualized detection of single events, we first provide a detailed cross-sectional characterization of age-specific patterns of slow and fast sleep spindles, slow oscillations, and their coupling in children aged 5 to 6, 8 to 11, and 14 to 18 years. Critically, based on this, we then investigated how spindle and slow oscillation maturity substantiate age differences in their precise orchestration. While the predominant type of fast spindles was development-specific in that it was still nested in a frequency range below the canonical fast spindle range for the majority of children, the well-known slow oscillation-spindle coupling pattern was evident for sleep spindles in the canonical (adult-like) fast spindle range in all three age groups – but notably less precise in children. To corroborate these findings, we linked personalized measures of fast spindle maturity, which indicate the similarity between the prevailing development-specific and adult-like fast spindles, and slow oscillation maturity, which reflects the extent to which slow oscillations show frontal dominance, with individual slow oscillation-spindle coupling patterns. Importantly, we found that fast spindle maturity was uniquely associated with enhanced slow oscillation-spindle coupling strength and precision. Taken together, our results suggest that the increasing ability to generate canonical fast sleep spindles actuates precise slow oscillation-spindle coupling patterns across child and adolescent development.

## 1 Introduction

The grouping of sleep spindles (SP, 9-16 Hz, Cox et al., 2017) into sequences of increased and decreased activity by the sleep slow oscillation (SO, 0.5-1 Hz, Steriade, 2006) during non-rapid eye movement sleep (NREM) has been recognized as an intrinsic property of the healthy, mature mammalian corticothalamic system for decades (Contreras et al., 1996; Contreras & Steriade, 1995; Mulle et al., 1986; Staresina et al., 2015; Steriade et al., 1993). The joint depolarization of large groups of cortical neurons during the SO up state impinges on neurons of the reticular thalamic nucleus, there, creating conditions that facilitate thalamic SP generation (Steriade, 1999). SPs then propagate to the cortex via thalamocortical projections, where they promote synaptic plasticity through changes in calcium activity (Niethard et al., 2018; Rosanova & Ulrich, 2005). However, it is still an open question how this precise coalescence develops across childhood and adolescence.

Far from being an epiphenomenon, accumulating evidence suggests that the synchronization of canonical fast SPs (≈ 12.5-16 Hz, centro-parietal predominance, Cox et al., 2017) precisely during the up state of SOs provides an essential mechanism for neural communication, e.g., supporting systems memory consolidation during sleep (Hahn et al., 2020; Helfrich et al., 2018; Mölle et al., 2002; Muehlroth et al., 2019). Importantly, canonical fast SPs in turn assort hippocampal ripples (Clemens et al., 2007; Helfrich et al., 2019; Siapas & Wilson, 1998; Staresina et al., 2015), that code for wake experiences and are considered a reliable marker of hippocampal memory consolidation (Buzsáki, 2015; Sirota et al., 2003). Moreover, canonical fast SPs themselves are associated with facilitated hippocampal-neocortical connectivity (Andrade et al., 2011; Cowan et al., 2020). In addition to canonical fast SPs, there is substantial evidence for a canonical slow SP type (≈ 9-12.5 Hz, frontal predominance, Cox et al., 2017) in human surface electroencephalogram (EEG, De Gennaro & Ferrara, 2003; Fernandez & Lüthi, 2020). However, previous findings hint at a distinct SO-slow SP coupling pattern and their function is still elusive (Klinzing et al., 2016; Mölle et al., 2011; Muehlroth et al., 2019; Rasch & Born, 2013). Taken together, the complex wave sequence of SO up state and canonical fast SPs, together with hippocampal activity, is considered to provide the scaffold for the precisely timed reactivation of initially fragile hippocampal memory representations and their transfer and integration into neocortical networks (Diekelmann & Born, 2010; Helfrich et al., 2019; Staresina et al., 2015). However, it is unclear whether the precise coupling of SP activity with SOs, described above, is already present or fully functional from early childhood on. Recent evidence indicates that the temporal co-ordination of SOs and SPs may improve across childhood and adolescence (Hahn et al., 2020; Joechner et al., 2021).

Likewise, the individual neural rhythms that define the coupling undergo substantial changes during child and adolescent development. Across maturation, SPs increase in occurrence and are found with a higher average frequency (Purcell et al., 2017; Zhang et al., 2021). Consistent with this, canonical slow SPs were reported to mature and dominate during early childhood. In contrast, canonical fast SPs are rarely detected in young children and become increasingly present and clearly dissociable only around puberty (D’Atri et al., 2018; Goldstone et al., 2019; Hoedlmoser et al., 2014; Shinomiya et al., 1999). However, amongst others, the application of individually adjusted frequency bands revealed that already young children express functional development-specific fast SPs in centro-parietal sites (D’Atri et al., 2018; Friedrich et al., 2019; Joechner et al., 2021; Zhang et al., 2021). Individualized rhythm detection methods provide an effective approach to capture true, dominant oscillatory rhythms despite substantial inter-individual variability, which presents a particular methodological challenge in developmental and age-comparative research (Cox et al., 2017; Muehlroth & Werkle-Bergner, 2020). While canonical fast SPs become more pronounced across development, an opposite trend can be observed for SOs (Buchmann et al., 2011; Kurth, Jenni, et al., 2010). Slow neuronal activity is initially maximally expressed and originates over posterior areas, developing towards a mature anterior predominance (Kurth, Ringli, et al., 2010; Timofeev et al., 2020). In summary, fast SPs and SOs separately are expressed differentially across development. However, it is still unclear how these developments interact to promote precise, adult-like temporal synchronization of SPs during SOs across childhood and adolescence.

Therefore, we aimed to (i) characterize the modulation of SPs during SOs across different ages and (ii) investigate how SP and SO maturity relate to the manifestation of SO-SP coupling. For this, we re-analyzed previously published nocturnal EEG data from a cross-sectional sample of 5-to 6-year-old children (Joechner et al., 2021) and a longitudinal cohort of children tested at 8 to 11 years of age (Hoedlmoser et al., 2014) and again at 14 to 18 years of age (Hahn et al., 2019, 2020).

## 2 Method

All analyses presented here are based on two previously published datasets: one cross-sectional cohort of 5-to 6-year-old children (Joechner et al., 2021) and one longitudinal sample of children, tested initially between ages 8 to 11 years (Hoedlmoser et al., 2014) and again around seven years later between ages 14 to 18 years (Hahn et al., 2019, 2020). Please refer to the original studies for a detailed description of all inclusion and exclusion criteria and the experimental procedures, respectively.

### 2.1 Participants

For the cross-sectional study, originally, 36 healthy children (19 female, *M*_age_ = 5 years, 9.53 months, *SD*_age_ = 6.50 months) were recruited from the database of the Max Planck Institute for Human Development (MPIB), Berlin, Germany, and from daycare centers in Berlin, Germany. Of these, five participants did not complete the study, and another seven children were excluded due to technical issues (*n* = 4) during polysomnography (PSG) or missing compliance with the study protocol (*n* = 3). Therefore, the same 24 5-to 6-year-old participants (13 female, *M*_age_ = 5 years, 10.71 months, *SD*_age_ = 7.28 months) as in Joechner et al. (2021) are presented here.

The initial sample of the longitudinal cohort consisted of 63 healthy children (T1, 28 female, *M*_age_ = 10 years, 1.17 months, *SD*_age_ = 7.97 months, Hoedlmoser et al., 2014), recruited from public elementary schools in Salzburg, Austria. Approximately seven years later, 36 participants returned for a follow-up assessment (T2, 24 female, *M*_age_ = 16 years, 4.56 months, *SD*_age_ = 8.76 months, Hahn et al., 2019). Two teens had to be excluded due to technical problems during PSG and data of one participant was not further analyzed because of insufficient amounts of non-rapid eye movement (NREM) sleep stage 3 (N3; out of four nights, in only two nights N3 sleep was detected; together: 3.05 %). Hence, we here present repeated-measures data for a total of 33 participants (23 female, *M*_ageT1_ = 9 years, 11.70 months, *SD*_ageT1_ = 8.35 months; *M*_ageT2_= 16 years, 4.91 months, *SD*_*a*geT2_ = 9.06 months). Participants and their families received a gift (5-to 6-year-old and 8-to 11-year-old participants) and/or monetary compensation (the parents of the 5-to 6-year-olds and the 14-to 18-year-olds) for their study participation. Both studies were designed in accordance with the Declaration of Helsinki and approved by the local ethics committee of either the MPIB, Germany, or the University of Salzburg, Austria. Given that participants were assessed during three distinct age intervals, the term “age group(s)” will be used to refer to 5-to 6-year-olds, 8-to 11-year-olds and 14-to 18-year-olds.

### 2.2 General procedure

Despite procedural differences, both studies comprised two nights of ambulatory PSG for each test time point. While the first night served to familiarize participants with the PSG, a memory task was performed before and after the second night (an associative scene-object task for the cross-sectional study and an associative word-pair task for both time points of the longitudinal study). Sleep was monitored in the habitual environment of each participant. For each 5-to 6-year-old participant, PSG recordings started and ended according to the individual bedtime. For the participants in the longitudinal sample, recordings were scheduled between 7:30-8:30 p.m. and 6:30 a.m. and were stopped after a maximum of 10 h time in bed during childhood. During the follow-up assessment, time in bed was fixed to 8 h between 11 p.m. to 7 a.m. Given well-known first night effects in children (Scholle et al., 2003), only the second night was analyzed here.

### 2.3 Sleep EEG acquisition and analyses

#### 2.3.1 Sleep recordings

For the youngest cohort, sleep was recorded using an ambulatory amplifier (SOMNOscreen plus, SOMNOmedics GmbH, Randersacker, Germany). A total of seven gold cup electrodes (Grass Technologies, Natus Europe GmbH, Planegg, Germany) were placed on the scalp for EEG recordings (F3, F4, C3, Cz, C4, Pz, Oz). EEG channels were recorded at a sampling rate of 128 Hz against the common online reference Cz. The signal of the AFz served as ground. Initial impedance values were kept below 6 kΩ. Additionally, two electrodes were placed at the left and right mastoids (A1, A2) for later re-referencing. Horizontal electrooculogram (EOG) was assessed bilaterally around the eyes and electromyogram (EMG) was recorded on the left and right musculus mentalis, referenced to one chin electrode. Furthermore, cardiac activity was monitored with two electrocardiogram (ECG) channels.

For the two older age groups, PSG signals were also collected ambulatory with a portable amplifier system (Varioport, Becker Meditec, Karlsruhe, Germany) at a sampling frequency of 512 Hz and against the common reference at Cz. For EEG recordings, 11 gold-plated electrodes (Grass Technologies, Natus Europe GmbH, Planegg, Germany; F3, Fz, F4, C3, Cz, C4, P3, Pz, P4, O1, O2) were placed along with A1 and A2 at the bilateral mastoids for offline re-referencing. Further, two horizontal and two vertical EOG channels, as well as two submental EMG channels, were recorded.

#### 2.3.2 Pre-processing and sleep staging

Sleep was staged automatically (Somnolyzer 24 × 7, Koninklijke Philips N.V.; Eindhoven, The Netherlands) and visually controlled by an expert scorer according to the criteria of the American Academy of Sleep Medicine (AASM; see Supplementary Table 1 for sleep architecture). Initial pre-processing was performed using BrainVision Analyzer 2.1 (Brain Products, Gilching, Germany). EEG channels were re-referenced offline against the average of A1 and A2 and filtered between 0.3-35 Hz. Further, all EEG data were resampled to 256 Hz. All subsequent analyses were conducted using Matlab R2016b (Mathworks Inc., Sherborn, MA) and the open-source toolbox Fieldtrip (Oostenveld et al., 2011). Firstly, bad EEG channels were rejected based on visual inspection. The remaining channels were then cleaned by applying an automatic artifact detection algorithm on 1-sec segments (see Joechner et al., 2021; Muehlroth et al., 2019 for more details).

#### 2.3.3 Sleep spindle and slow oscillation detection

SPs were detected during artifact-free NREM (N2 and N3) epochs at frontal (F3, F4) and centro-parietal channels (C3, C4, Cz, Pz) using an established algorithm (Klinzing et al., 2016; Mölle et al., 2011; Muehlroth et al., 2019) with individual amplitude thresholds. Frequency windows for SP detection were defined based on two approaches: (i) An individualized approach, aiming at capturing the person- and development-specific dominant rhythm, and (ii) a fixed approach, targeting “adult-like” fast SPs in the canonical fast SP range that were shown to be coupled to SOs before, despite missing evidence for their strong presence in pre-school children (Joechner et al., 2021, see Figure 1A and Figure 3). For the individualized approach, we defined the frequency bands of interest as the participant-specific peak frequency in the averaged frontal (F3, F4) or centro-parietal (C3, Cz, C4, Pz) background-corrected NREM power spectra (9-16 Hz) ± 1.5 Hz (see Figure 1 for peak distributions; Mölle et al., 2011; Ujma et al., 2015). Therefore, NREM power spectra were calculated within participants for averaged frontal and centro-parietal electrodes between 9-16 Hz by applying a Fast Fourier Transform (FFT) and using a Hanning taper on 5-sec epochs. To restrict the search space for peak detection to dominant rhythmic activity (Aru et al., 2015; Kosciessa et al., 2020), we then modeled the background spectrum and subtracted it from the original power spectrum. Assuming that the EEG background spectrum follows an A*f^-a^ distribution (Buzsáki & Mizuseki, 2014; He et al., 2010), background spectra were estimated by fitting the original power spectrum linearly in the log(power)-log(frequency) space employing robust regression (Kosciessa et al., 2020). Frontal and centro-parietal peaks were finally detected in the background-corrected power spectra by a classical search for maxima combined with a first derivative approach (Grandy et al., 2013). For the fixed approach, frequency bands were restricted to 12.5-16 Hz in every individual – representing the canonical range for fast SP extraction in adults – and detection was focused on averaged centro-parietal electrodes only to capture adult-like fast SPs (C3, Cz, C4, Pz; Cox et al., 2017). For detection of SP events, EEG data were first filtered in the respective frequency bands using a 6^th^ order two-pass Butterworth filter (separately for individually identified frontal and centro-parietal, as well as adult-like fast centro-parietal SPs). Then, the root mean square (RMS) was calculated using a moving window of 0.2 sec and smoothed with a moving average of 0.2 sec. Finally, a SP was detected whenever the envelope of the smoothed RMS signal exceeded its mean by 1.5 *SD* of the filtered signal for 0.5-3 sec. SPs with boundaries within a distance between 0-0.25 sec were merged as long as the resulting event was shorter than 3 sec (see e.g., Mölle et al., 2011; Muehlroth et al., 2019 for similar methods; see Supplementary Figures 1 and 2 for an example of how individually identified and adult-like fast centro-parietal SPs manifested in the raw EEG and how the average shape of these SPs looked like in one 6-year-old and one child tested at 9 and again at 15 years of age).

**Figure 1.**
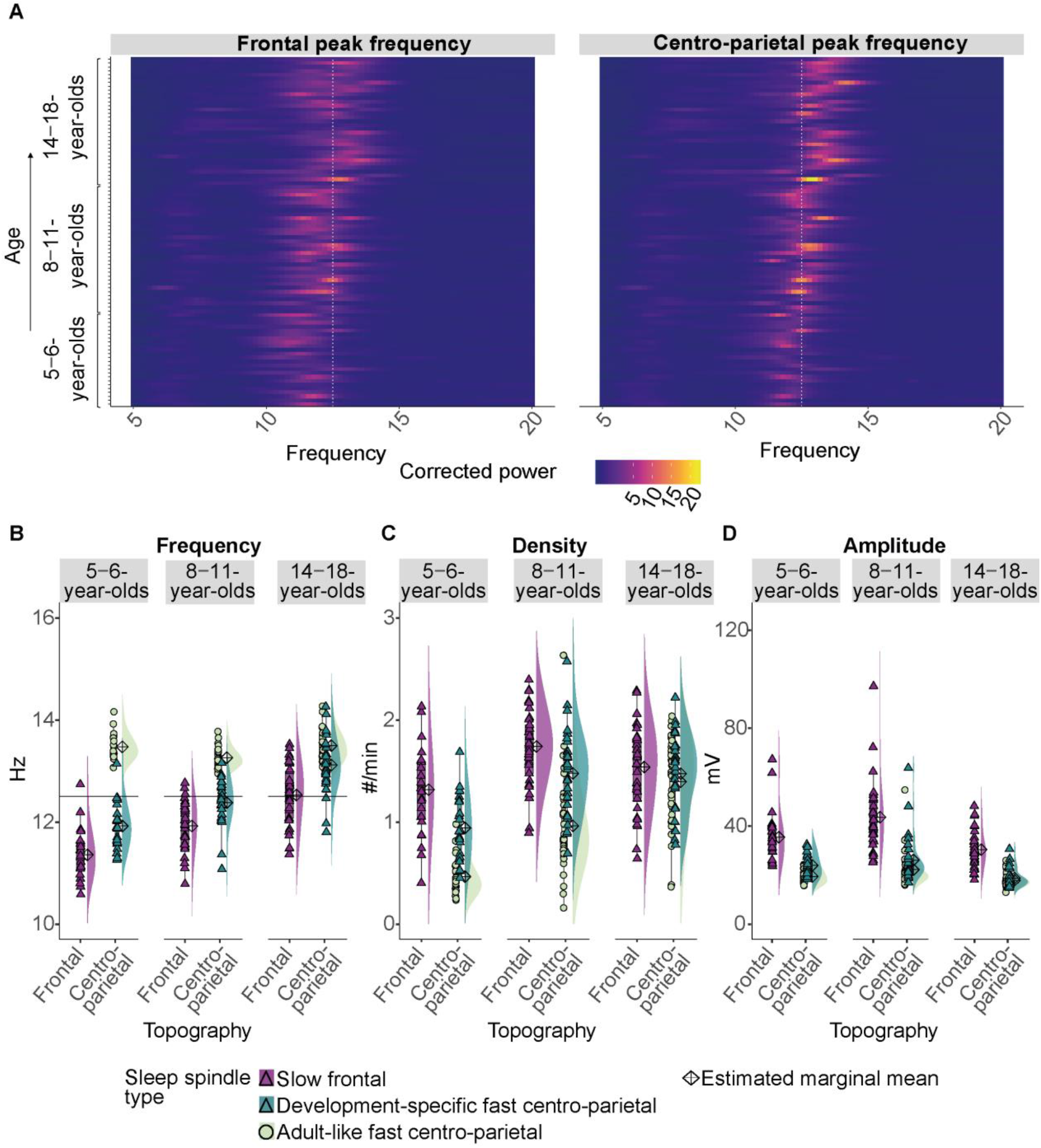
(A) Background-corrected power spectra in averaged frontal (left) and averaged centro-parietal (right) electrodes showing thepeak frequencyin thesleep spindlefrequency range(9 −16 Hz) in brightercolors forevery participant at every age. Data are ordered by age from bottom to top. The white dotted line at 12.5 Hz illustrates the frequency border for canonical fast sleep spindles. Note: Children at an older age (bottom to top) showed higherpeak frequencies. Further, a linear mixed-effects model revealed that peak frequencies in centro-parietal electrodes were significantly faster compared to frontal ones within all age groups. However, formost of the young children (lowerpart of the plot) the peak frequencies and the frequencies of the derived dominant fast spindles (B) were slower than the canonical band (development-specific) (B-D) Results from the comparison of adult-like fast (12.5-16 Hz) and individually-identified slow frontal and development-specific fast centro-parietal sleep spindle (B) frequency, (C) density, and (D) amplitude for all three age groups. Values are individual raw scores. Diamonds reflect estimated marginal means of the respective linearmixed-effects model.

**Figure 2.**
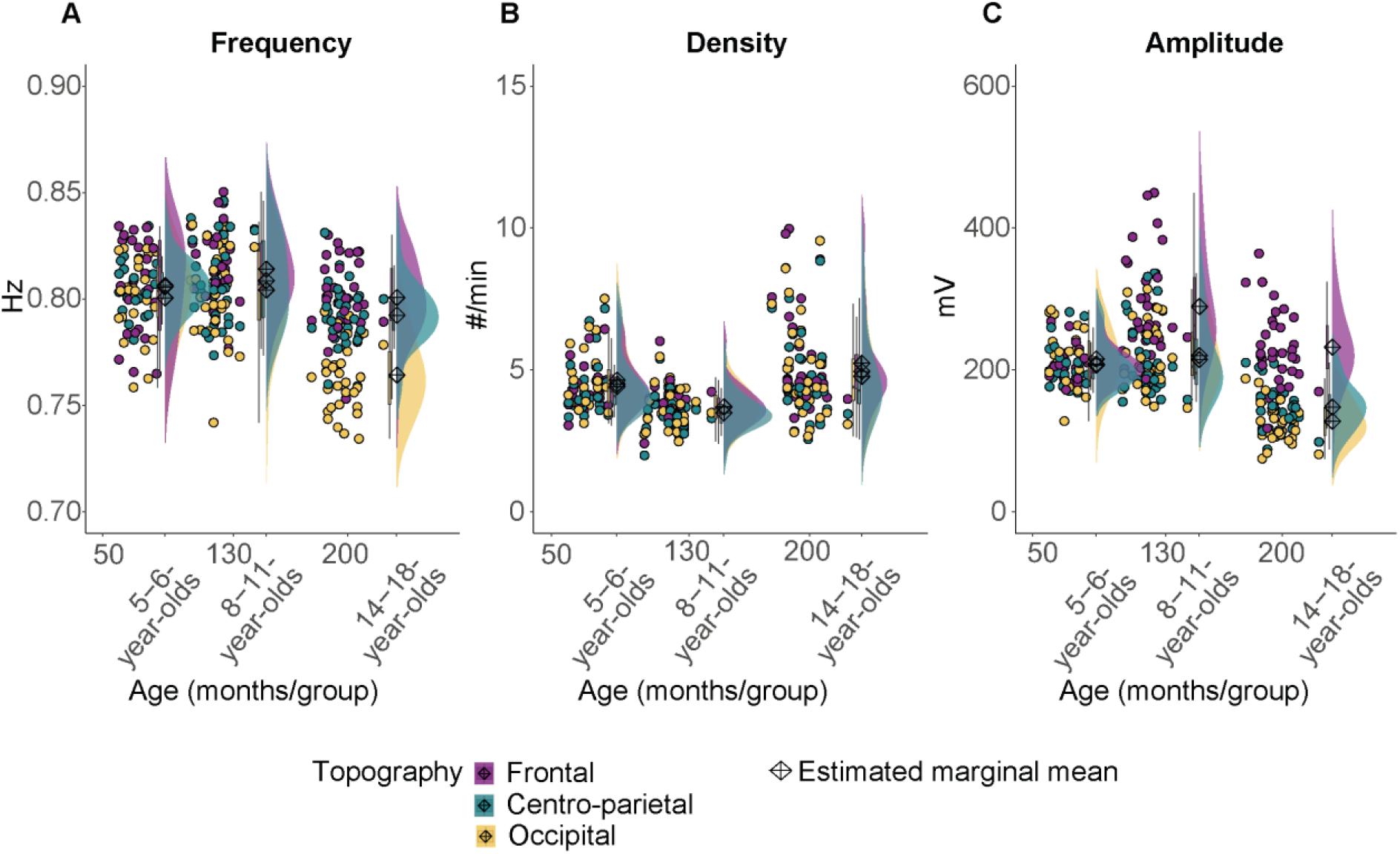
Results from the comparison of slow oscillation (A) frequency, (B) density, and(C) amplitude forall three age groups. Values are individualraw scores. Diamonds reflect estimated marginal means of the respective linear mixed-effects model. Note that we observed the strongest age-related differences for the slow oscillation amplitudein different topographic locations.

**Figure 3.**
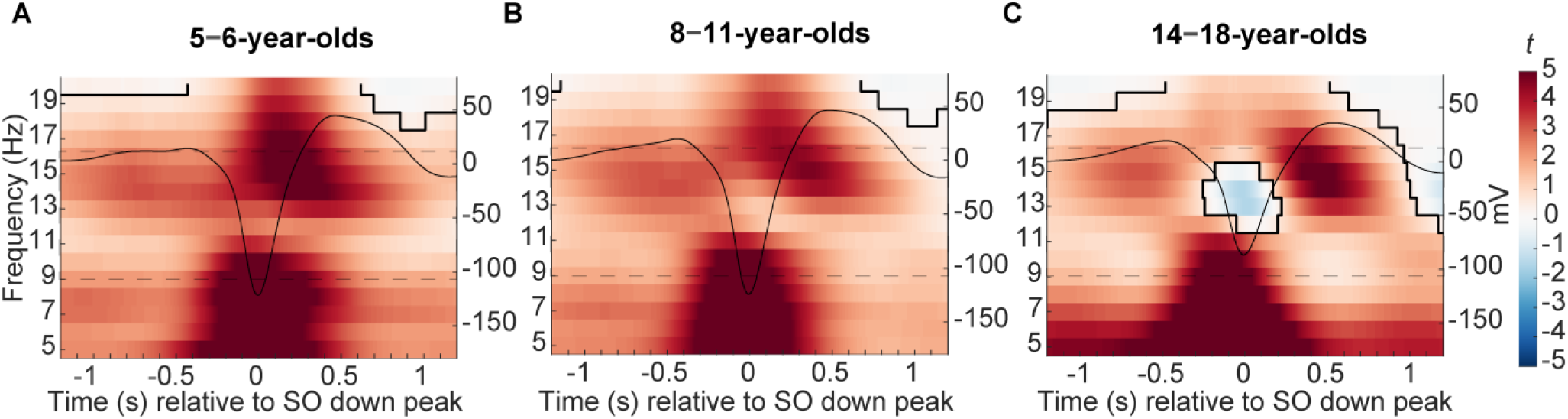
Centro-parietal power differences between trials with and without centro-parietal slowoscillations (in *t*-score units). Significant clusters are outlined in black (cluster-based permutation test, cluster α < .05, two-sided test). The average centro-parietal slow oscillation foreach agegroup (A-C) is plotted onto thepower differences in black to illustrate the relation to slow oscillation phase(scalein mV on the right y-axis of each plot). The sleep spindle frequency range is highlighted by the dashed window. Note that the strongest power increases during slow oscillations were observed ina frequency range reflecting the canonical fast sleep spindle range (12.5-16 Hz). Results for frontal powerand frontal slow oscillations can be found in Supplementary Figure 5.

SO detection was based on an algorithm with individually adapted amplitude thresholds (Mölle et al., 2002; Muehlroth et al., 2019) and performed for all NREM epochs. First, the EEG signal was filtered between 0.2 and 4 Hz using a two-pass Butterworth filter of 6^th^ order. Subsequently, zero-crossings were marked to identify positive and negative half-waves. Pairs of negative and succeeding positive half-waves were considered a potential SO if their frequency was 0.5-1 Hz. Only putative SOs with a peak-to-peak amplitude of 1.25 times the mean peak-to-peak amplitude of all tagged SOs and a negative amplitude of 1.25 times the average negative amplitude of all SOs that did not include artifact segments were kept for the following analyses.

#### 2.3.4 Temporal association between sleep spindles and slow oscillations

To assess the temporal association between SPs and SOs, we employed two different approaches following previous reports in developmental samples (Joechner et al., 2021; Muehlroth et al., 2019).

##### 2.3.4.1 Time-frequency analyses

On the one hand, we examined power modulations during SOs between 5-20 Hz within intervals of ± 1.2 sec around the down peak of SOs. For this, we detected artifact-free NREM epochs containing SOs (down peak ± 3 sec) and equally long, randomly chosen segments without SOs. Time-frequency representations (Figure 3) were then calculated for trials with and without SOs using a Morlet wavelet transformation (12 cycles) in steps of 1 Hz and 0.002 sec. The resulting time-frequency pattern of trials with and without SOs was contrasted for every participant using independent sample *t*-tests. Subsequent group-level analyses were conducted separately for the cross-sectional cohort and the two cohorts from the longitudinal sample. Within these three age groups, the *t*-maps were then contrasted against zero using a cluster-based permutation test (two-sided, critical alpha-level α = .05; note for the ease of comparability, *p*-values for all cluster-based permutation tests were multiplied by 2 and thus values below .05 were considered significant; Maris & Oostenveld, 2007) with 5,000 permutations in a time window between −1.2 to +1.2 sec around the SO down peak.

##### 2.3.4.2 Temporal co-occurrence of sleep spindle events with slow oscillations

On the other hand, we investigated the temporal co-occurrence of individually identified and adult-like fast SP events with SOs. In a first step, we calculated the general co-occurrence rate between SPs and SOs on a broad time scale by determining the percentage of SP event centers that occurred within an interval of ± 1.2 sec around SO down peaks during NREM sleep, relative to all SPs during NREM sleep. We also repeated this analysis vice versa for SO down peaks relative to SP center occurrence (see Supplementary Figures 3-4 and Supplementary Tables 2-7).

To specify the temporal relation between SP and SO co-occurrence on a finer temporal scale, in a second step, we calculated peri-event time histograms (PETH, Figure 4). Therefore, the intervals of ± 1.2 sec around the SO down peak were partitioned into 100 ms bins and the proportion of SP centers occurring within each time bin was assessed. The occurrence rates within a bin were subsequently normalized by the total counts of SPs occurring during the complete respective SO down peak ±1.2 sec interval and multiplied by 100. To determine whether SP activity was modulated during SOs differently from what would be expected by chance, we created a surrogate comparison distribution for every participant during every test time point. The original percentage frequency distribution was randomly shuffled 1,000 times and averaged across all permutations. The resulting surrogate distributions were then compared against the original distributions using dependent sample *t*-tests. To account for the multiple comparisons, a cluster-based permutation test with 5,000 permutations was implemented (two-sided, critical alpha-level α = .05, *p*-values for all cluster-based permutation tests were multiplied by 2 and thus values below .05 were considered significant; Maris & Oostenveld, 2007). All analyses were conducted separately for individually identified SPs at frontal (averaged over F3, F4) and centro-parietal (averaged over C3, Cz, C4, Pz) electrodes and for adult-like fast SPs detected at centro-parietal (averaged over C3, Cz, C4, Pz) derivations.

**Figure 4.**
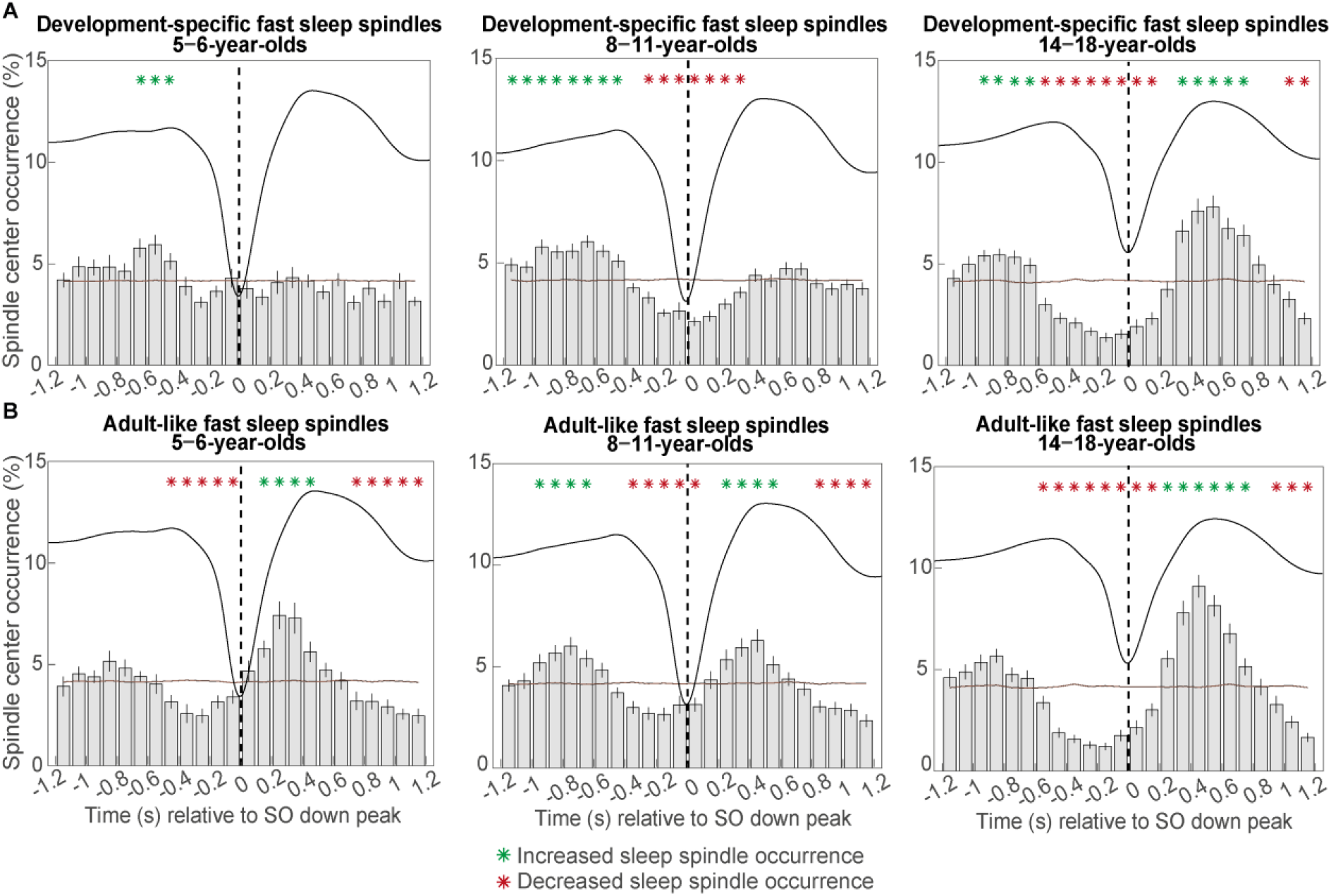
Peri-event time histograms for (A) development-specific fast centro-parietal spindles and (B) adult-like fast centro-parietal spindles showing theproportionof events occurring within 100 ms bins during centro-parietal slow oscillations. Errorbars represent standard errors. Green asterisks mark increased spindle occurrence (positive cluster, cluster α < .05, two-sided test) and red asterisks mark decreased spindle occurrence (negative cluster, clusterα <.05, two-sided test) compared to random occurrence(horizontal line). Thedashedverticallineindicates the slow oscillation down peak. Theaveragecentro-parietal slow oscillation of each agegroup is shown in black to illustrate the relation to the slow oscillation phase. Results for slow oscillations and spindles in different topographies can be foundin Supplementary Figure 6.

To quantify the modulation strength of SP occurrence during SOs, for every participant, we calculated the Kullback-Leibler (KL) divergence between the percentage frequency distribution (p) and its surrogate (q) used for the PETHs. The KL divergence is a measure rooted in information theory that describes the amount of information loss if one distribution was approximated by the other (Joyce, 2011). This measure is the basis for several commonly used methods to determine phase-amplitude coupling and was calculated in the following way:

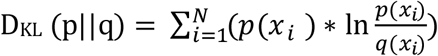

The higher the value, the more two distributions deviate from each other. Hence, higher values indicate that the actual percentage frequency distribution of the PETHs deviated from a uniform distribution, i.e., the more likely SPs were concentrated around specific phases of the underlying SO.

### 2.4 Statistical analyses

#### 2.4.1 Age differences in sleep spindles and slow oscillations

Age-related comparisons between SP and SO parameters were conducted using linear mixed-effects models (LMMs) with restricted maximum likelihood variance estimation (REML) and the bobyqa optimizer from the NLopt library (Powell, 2009). LMMs were implemented in R 4.0.3 (R Core Team, 2020) using Rstudio Version 1.1.383 and the lme4 package (Bates et al., 2015). Given that our data consist of both, purely cross-sectional as well as repeated measures, we set up the combined data set mimicking a longitudinal design with a total of 57 (*n* cross-sectional + *n* longitudinal) participants, each tested three times between 5 to 6, 8 to 11, and 14 to 18 years of age. While for the 5-to 6-year-olds, the 33 participants from the longitudinal cohort were assigned missing values, for the 8-to 11- and 14-to 18-year-olds, the 24 participants of the cross-sectional sample were treated as missing. It is well established that LMMs perform well even with a larger number of missing data (Cnaan et al., 1997; Krueger & Tian, 2004). Fixed effects were most of the time “age group” (i.e., 5-to 6-, 8-to 11-, 14-to 18-year-olds) and “topography” (e.g., frontal, central, occipital) and specified based on our research questions. The effects of categorical predictors were set up as sum-coded factors. To account for the nonindependence when dealing with repeated measures, participants were included as random effects. Overall, only models with a random intercept for participants converged (but not with a random slope). Post-hoc pairwise comparisons were conducted using the emmeans package (Lenth, 2021). Degrees of freedom (*df*) were calculated applying the Satterthwaite method (Giesbrecht & Burns, 1985; Kuznetsova et al., 2017) and *p*-values were Bonferroni corrected (*p*_adj_, Bland & Altman, 1995).

#### 2.4.2 Association between sleep spindle and slow oscillation maturity with slow oscillatio n-spindle coupling

To assess the association between SP and SO maturity and SO-SP coupling (for an exact definition of the variables, see the results section), we conducted two analyses:

First, a partial least squares correlation analysis (PLSC, Krishnan et al., 2011) was calculated between indicators of SP amplitude, frequency, and density maturity and the *t*-maps of power modulations during SOs (compared to non-SO trials, time-frequency maps). PLS methods are multivariate tools to extract commonalities between two data sets using singular value decomposition. PLSC specifically analyzes the association between two data sets by deriving pairs of singular vectors that cover the maximal covariance between two matrices (Krishnan et al., 2011). Singular vector pairs are ordered by the amount of covariance they contribute to the association as reflected by their corresponding singular values. The components of the singular vectors represent weights (also called saliences) that define how each element of the data sets contributes to a given association. Ultimately, the projection of pairs of saliences onto their original data matrices results in pairs of latent variables that capture the maximal amount of common information between the two data sets (e.g., in our case a latent time-frequency and latent SP maturity variable). Therefore, PLSC is ideally suited to identify time-frequency patterns associated with a specific SP maturity profile. The number of impactful singular vector pairs (i.e., those that account for a significant amount of covariance) was identified by a permutation test with 5,000 permutations based on the singular values. For each significant vector pair, reliability of the weights of each time-frequency value, defining the SO-SP coupling pattern associated with a SP maturity pattern, was determined using bootstrap ratios (BSR). BSRs represent the ratio of the weights of each time-frequency value and their bootstrap standard errors based on 5,000 samples. BSRs are akin to *Z*-scores, hence BSRs higher than 1.96 and lower than −1.96 were considered stable (Krishnan et al., 2011). The SP maturity profile was represented as the correlation between the latent time-frequency variable and the three raw SP maturity variables. These values are comparable to the SP maturity weights and indicate whether the association between the time-frequency pattern and individual SP maturity variables is in the same or different direction (McIntosh & Lobaugh, 2004). Stability of these correlation patterns was determined by their bootstrap estimated 95% confidence intervals (McIntosh & Lobaugh, 2004). Note, despite our awareness of different dependencies between the age groups, 5-to 6-, 8-to 11-, and 14-to 18-year-olds were considered for this analysis.

Second, generalized LMMs (GLMM) were used to examine the relation between the modulation of individually identified SPs (as captured by the KL divergence of the PETHs) with SP and SO maturity. We defined the first component of a principal component analysis (PCA) between indicators of SP amplitude, frequency, and density maturity across the complete data set (across all age groups) as a general SP maturity factor. We then aimed to associate the KL divergence values reflecting individually identified SP modulation strength during SOs with the general SP maturity component. For this, we set up log-linked gamma GLMMs with the KL divergence as the dependent variable and with the SP maturity component as a fixed factor, allowing for random intercepts per participant. In addition, an indicator of SO maturity was added as a covariate. We used the bobyqa optimizer from the minqa package (Bates et al., 2014).

### 3 Results

### 3.1 Development-specific fast centro-parietal sleep spindles become more prevalent and increasingly resemble adult-like fast sleep spindles with older age

In the first step, we aimed at identifying age-specific patterns of SPs. Using LMMs, we first contrasted the peak frequencies of the power spectra at frontal and centro-parietal sites between age groups and topographies (frontal, centro-parietal) to obtain evidence for two distinct oscillatory rhythms (Figure 1A).

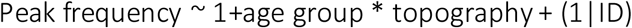

Similarly, we then compared features of individually identified SPs (i.e., frequency, density, amplitude; Figure 1B-D), which were detected based on the peak frequencies in frontal or centro-parietal sites, and thus represent the dominant SP rhythms per individual.

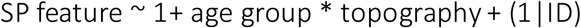

For both, spectral SP peak frequency and the average frequency of SP events, we found significant main but no interaction effects (analysis of variance (ANOVA) outputs, Supplementary Tables 8 and 10). Follow-up pairwise comparisons revealed that peak frequency and the average frequency of SP events, averaged across age groups, were significantly higher for centro-parietal electrodes compared to frontal sites (peak frequency: *t*(120.05)= −8.31, *p*_*adj*_ < .001; average SP event frequency: *t*(119.51) = −11.85, *p*_*adj*_ < .001). Further, SP frequency averaged across topographical locations was higher for 14-to 18-year-old compared to 8-to 11-year-old compared to 5-to 6-year-old participants (all *t* ≤ −3.85 all *p*_*adj*_ ≤ .001; for pairwise comparison results see Supplementary Tables 9 and 11).

Hence, across both measures and age groups individually identified, dominant frontal SPs revealed lower frequencies than centro-parietal SPs, indicating the presence of two distinguishably fast SP types. We will henceforth be referring to slow frontal and fast centro-parietal SPs, respectively. Crucially, despite being faster, the frequency of the fast centro-parietal SPs was specific to the age of the participants in a way, that at a younger age, these SPs were not yet in the range of canonical fast SPs. Instead, they were nested in the canonical slow SP range. Hence, the term “development-specific” will be added for individually determined fast centro-parietal SPs.

For density and amplitude, all simple effects and the interaction between age group and SP topography reached significance (Supplementary Tables 12 and 14). Pairwise comparisons indicated higher density for slow frontal compared to development-specific fast centro-parietal SPs for the 5-to 6-(*t*(119.22) = 5.54, *p*_*adj*_ < .001) and the 8-to 11-year-olds(*t*(119.22) = 4.50, *p*_*adj*_ < .001), while the difference was not significant for 14-18-year-olds (*t*(119.22)= 0.81, *p*_*adj*_ = 1.000). The interaction for SP amplitude was mainly driven by a significantly higher slow frontal amplitude for the 8-to 11-year-olds compared to the 5-to 6-(*t*(103.18)= −3.39, *p*_*adj*_ = .015) and the 14-to 18-year-olds (*t*(120.25) = 8.94, *p*_*adj*_ < .001) and a lower development-specific fast centro-parietal amplitude for the 14-to 18-compared to the 8-to 11-year-olds (*t*(120.25) = 4.99, *p*_*adj*_ < .001). Further within all age groups amplitudes were lower for development-specific fast compared to slow frontal SPs (all *t* ≥ 6.66, *p*_ad*j*_ < .001; see Supplementary Tables 13 and 15 for all post-hoc comparison results).

To sum up, ourdata indicate theexistence of slow frontal and development-specific fast centro-parietal SPs at all ages under study. Specifically, the development-specific fast centro-parietal SPs become increasingly expressed and faster with older age. Given evidence for the specific role of canonical fast SPs for memory (Rasch & Born, 2013), henceforth, we will focus on SPs detected at centro-parietal electrodes (however, corresponding analyses were also conducted for slow frontal SPs and can be found in the Supplementary Material).

After having characterized development-specific fast centro-parietal SPs across all our age groups, we were interested in how they differed from canonical fast SPs. Therefore, we additionally extracted adult-like fast SPs in centro-parietal electrodes by applying fixed frequency criteria between 12.5-16 Hz – the frequency band commonly referred to as fast in adults (Cox et al., 2017; see Supplementary Figure 1 for an example raw EEG trace with development-specific and adult-like fast SPs and Supplementary Figure 2 for examples of average detected events). We then compared characteristics (i.e., frequency, density, amplitude; Figure 1B-D) of development-specific fast centro-parietal SPs with adult-like fast centro-parietal SPs using LMMs across the three age groups. In addition to “age group”, the factor “SP type” (development-specific vs. adult-like) was entered as a fixed effect:

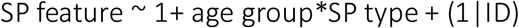

Results indicated that all main effects and the interaction between the factors “age group” and “SP type” were significant for frequency, density, and amplitude (for a summary of all analyses and all post-hoccomparisons see Supplementary Tables 16-21). Post-hocpairwisecomparisons revealed that the frequency of adult-like fast SPs was consistently higher as compared to the development-specific fast centro-parietal SPs within all age groups (*t*_*5-6*_ (124.05)= 18.19, *p*_*adj*_ < .001; *t*_*8-11*_(124.05) = 12.13, *p*_*adj*_ < .001; *t*_*14-18*_ (124.05) = 5.22, *p*_*adj*_ < .001). In line with a generally higher frequency, in 5-to 6- and 8-to 11-year-olds, adult-like fast SPs had a lower amplitude compared to development-specific fast centro-parietal SPs (*t*_*5-6*_ (122.53)= −4.06, *p*_*adj*_ = .001; *t*_*8-11*_(122.53) = −4.18, *p*_*adj*_ < .001). Crucially, this effect was non-significant in the oldest age group (*t*_*14-18*_ (122.53)= −0.92, *p*_*adj*_ = 1.000). Similarly, whereas density of development-specific fast SPs was higher compared to adult-like fast SP density in the 5-to 6- and the 8-to 11-year-olds (*t*_*5-6*_ (122.60)= −7.23, *p*_*adj*_ < .001; *t*_*8-11*_(122.60) = −9.14, *p*_*adj*_ < .001), there was no difference in density for the 14-to 18-year-olds (*t*_*14-18*_ (122.60)= −1.41, *p*_*adj*_ = 1.000). Indeed, density of adult-like fast SPs was highest for the oldest age group compared to the 8 - to 11-(*t*(122.60) = −7.67, *p*_*adj*_ < .001) and 5-to 6-year-olds (*t*(97.35)= −9.38, *p*_*adj*_ <.001) and higher for the 8-to 11- as compared to the 5- to 6-year-olds (*t*(97.35) = −5.02, *p*_*adj*_ < .001). Density for the development-specific fast centro-parietal SPs was only lower for the 5- to 6-year-olds as compared to both older age groups (*t*_*8-11*_ (97.35) = −5.40, *p*_*adj*_ < .001; *t*_*14-18*_(97.35) = −5.36, *p*_*adj*_ < .001) while there was no difference between the 8- to 11- and 14- to 18-year-olds (*t*(122.60) = 0.06, *p*_*adj*_ =1.000).

Crucially, this indicates that despite the absence of a prominent peak in the power spectrum (Figure 1A), already young children express adult-like fast SPs which occur increasingly often with older age. Despite a higher frequency of the adult-like fast SPs in all age groups, with older age, the differences in amplitude and density between development-specific and adult-like fast centro-parietal SPs decrease and are no more evident in the oldest age group. Hence, it appears as if fast centro-parietal SPs become more dominant and adult-like in their frequency and amplitude characteristics in older children.

### 3.2 Slow oscillations dominate anteriorly only in older children and adolescents

Similar to the analyses on SPs, we next compared features that define SOs (i.e., frequency, density, amplitude; Figure 2) between age groups and between SOs detected at frontal (averaged over F3, F4), centro-parietal (averaged over Cz, Pz), and occipital (cross-sectional sample: Oz, longitudinal sample: averaged over O1, O2) electrodes (topography factor) using LMMs:

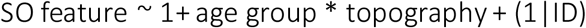

Occipital locations were added based on observations suggesting a posterior dominance of SOs in young children (Kurth, Ringli, et al., 2010; Timofeev et al., 2020). Further, considering evidence suggesting that surface SOs might be most powerful medially (Murphy et al., 2009), we focused on midline electrodes, while also keeping the overlap of electrodes between our samples as high as possible.

Results revealed that the frequency of SOs differed over topographic locations across age groups (age group*topography interaction: *F*(4,207.33) = 11.68, *p* < .001; see also Supplementary Table 22). Pairwise comparisons (see Supplementary Table 23 for all pairwise comparisons) indicated that SO frequency differed between recording sites only for the 14- to 18-year-olds. Both, frontal and centro-parietal frequency was higher compared to occipital frequency (*t*_*frontal-occipital*_ (207.33) = 10.25, *p*_*adj*_ < .001; *t*_*centro-parietal-occipital*_ (207.33) = 7.91, *p*_*adj*_ < .001). For SO density, the ANOVA output indicated a significant simple effect of age group (Supplementary Table 24, *F*(2,89.22) = 81.82, *p* < .001) but neither for topography nor the interaction effect. Pairwise comparisons revealed that density, averaged across topographical recording sites, was lower for the 8- to 11- compared to both, the 5- to 6- (*t*(61.98) = 3.25, *p*_*adj*_ = .006) and the 14- to 18-year-olds (*t*(206.49)= −12.77, *p*_*adj*_ < .001; see Supplementary Table 25 for all pairwise comparisons). For SO amplitude, inspection of the ANOVA output and the pairwise comparisons revealed that only the 8- to 11-year-olds and adolescents expressed SOs at a higher amplitude at frontal as compared to both centro-parietal and occipital locations (8- to 11-year-olds: *t*_*frontal-centro-parietal*_(208.07) = 10.89, *p*_*adj*_ < .001; *t*_*frontal-occipital*_(208.07) = 10.06, *p*_*adj*_ < .001; 14- to 18-year-olds: *t*_*frontal-centro-parietal*_(208.07) = 12.35, *p*_*adj*_ < .001; *t*_*frontal-occipital*_(208.07) =15.24, *p*_*adj*_ < .001; see Supplementary Tables 26-27 for all results).

To summarize, we did not observe a frontal prevalence of SOs in the youngest children. However, this dominance is present in older children and adolescents. Therefore, similar to our SP analyses, we will concentrate on centro-parietal SOs for all subsequent analyses. Results for SPs and SOs in additional topographical locations can be found in the Supplementary Material.

### 3.3 Temporal modulation of sleep spindle power during slow oscillations differs with age

We observed robust age differences in SPs and SOs across the three age groups. But do they also affect the temporal coupling between these two neural rhythms? In the first step, we aimed at determining the spectral and temporal characteristics of SO-coupled SPs. Hence, we compared spectral power (5-20 Hz) over centro-parietal recording locations during trials with and without centro-parietal SOs (± 1.2 Hz around down peak). Within all age groups, power in a broad range including the SP frequency band (9-16 Hz) was significantly higher during SOs as compared to trials without SOs (all cluster *p*s < .001) suggesting temporal clustering of SPs during the SO cycle. On a descriptive level (Figure 3), the strongest power differences during SOs as compared to trials without SOs for all three age groups were located within the frequency range of canonical fast SPs (12.5-16 Hz) within one second after the down peak, close to the up peak. However, the modulation of power in the canonical fast SP frequency range seemed to be stronger and more precise in our oldest age group. Specifically, for the younger children, the increased power in the canonical fast SP frequency range was surprising, given the overall lower density and power of adult-like fast SPs in children (Figure 1).

### 3.4 How does the temporal modulation of development-specific fast and adult-like fast sleep spindles during slow oscillations differ with age?

Having identified evidence for the temporally ordered occurrence of SPs during specific SO phases in all our age groups, we were interested in the next step in the precise temporal co-ordination of SP events and SOs. Therefore, we created PETHs for development-specific fast SPs. We compared the resulting occurrence-percentage distribution per participant with a participant-specific surrogate distribution to identify patterns of increased and decreased SP occurrence during the SO cycle.

As can be inferred from Figure 4A, we observed a shift towards a clear coupling from the youngest to the oldest age group. Only the oldest age group reliably presented the canonical coupling pattern known from adults: A decreased SP probability during the SO down state and an increased occurrence during the up state. Statistically, we found one positive cluster (*p* = .001) for the 5- to 6-year-olds from −700 to −400 ms for the modulation of development-specific fast centro-parietal SPs (up peak at ≈ 465 ms). For the 8- to 11-year-olds, we identified a more extended positive cluster from −1,200 to −400 (*p* < .001) and one negative cluster from - 300 to 400 sec (*p* < .001) for development-specific fast centro-parietal SPs (up peak at ≈ 488 ms). Lastly, for the 14- to 18-year-olds, PETHs indicated two positive clusters. One from 300 to 800 ms (*p* = .002) and another from −1,000 to - 600 ms (*p* = .014) and two negative clusters from - 600 to 200 ms (*p* < .001) and from 1,000 to 1,200 (*p* = .034) for development-specific fast centro-parietal SPs (up peak at ≈ 532 ms).

Critically, and somewhat unexpectedly for the two younger age groups, the previous analysis of spectral power differences in the SP frequency range along the SO cycle suggested temporal SO-SP alignment specifically for events in the canonical fast SP range. Hence, we repeated the PETH analyses for adult-like fast SPs. As can be inferred from Figure 4B, we observed a clear coupling of SP occurrence rates to specific phases of the SO cycle within all age groups – though less precise in the younger age groups.

For 5- to 6-year-olds, we identified one positive cluster (*p* < .001) from 100 to 500 ms and two negative clusters (both *p*s < .001) from −500 to 0 ms and from 700 to 1,200 ms for adult-like fast centro-parietal SPs (up peak at ≈ 465 ms). For the 8- to 11-year-olds, we found two positive clusters from −1,000 to −600 ms (*p* < .001) and from 200 to 600 ms (*p* = .001) and two negative clusters (both *p*s < .001) from −400 to 100 ms and from 800 to 1,200 ms (up peak at ≈ 488 ms). Lastly, for the 14- to 18-year-olds, we identified one positive cluster (*p* < .001) from 200 to 800 ms and two negative clusters from −600 to 200 ms (*p* < .001) and from 900 to 1,200 ms (*p* = .004) for adult-like fast centro-parietal SPs (up peak at ≈ 532 ms).

To summarize a clear coupling of SP occurrence at specific phases of the SO can hardly be detected for development-specific fast centro-parietal SPs in the youngest age group. Importantly, adult-like fast SPs are already modulated by SOs in the youngest age group – even though they only occur very rarely. However, independent of the SP type, the coalescence of SPs and the up peak of SOs becomes more precise with older age. Hence, despite clearly identifiable development-specific fast SPs, precise SO-SP coupling seems to depend on the presence of adult-like fast SPs.

### 3.5 Association between slow oscillation-spindle coupling and sleep spindle and slow oscillation maturity

So far, the analyses suggested that SPs in the canonical fast frequency range rather than the more dominant development-specific fast SPs are coupled to SOs. Hence, we reasoned, that the maturation of development-specific fast SPs towards adult-like fast SPs may explain age-related differences in the strength of the SO-SP coupling.

To capture the “maturational stage” of fast SPs, we computed the distance between development-specific and adult-like fast centro-parietal SPs within each participant. Specifically, we calculated difference measures in SP characteristics (i.e., frequency, density, amplitude) between adult-like and development-specific fast SPs. Note, given the opposing signs of the fast SP maturity differences, we inverted the frequency difference values. Further, we re-scaled the frequency, density, and amplitude difference scores using *Z*-transformation across all participants to convert all metrics into a common space. As illustrated in Figure 5A-C, the increasing *Z*-transformed difference scores (fast SP maturity scores) with older age of the participants capture the fact that the dominant, i.e., development-specific fast centro-parietal, SPs resemble more and more adult-like fast SPs in older children.

**Figure 5.**
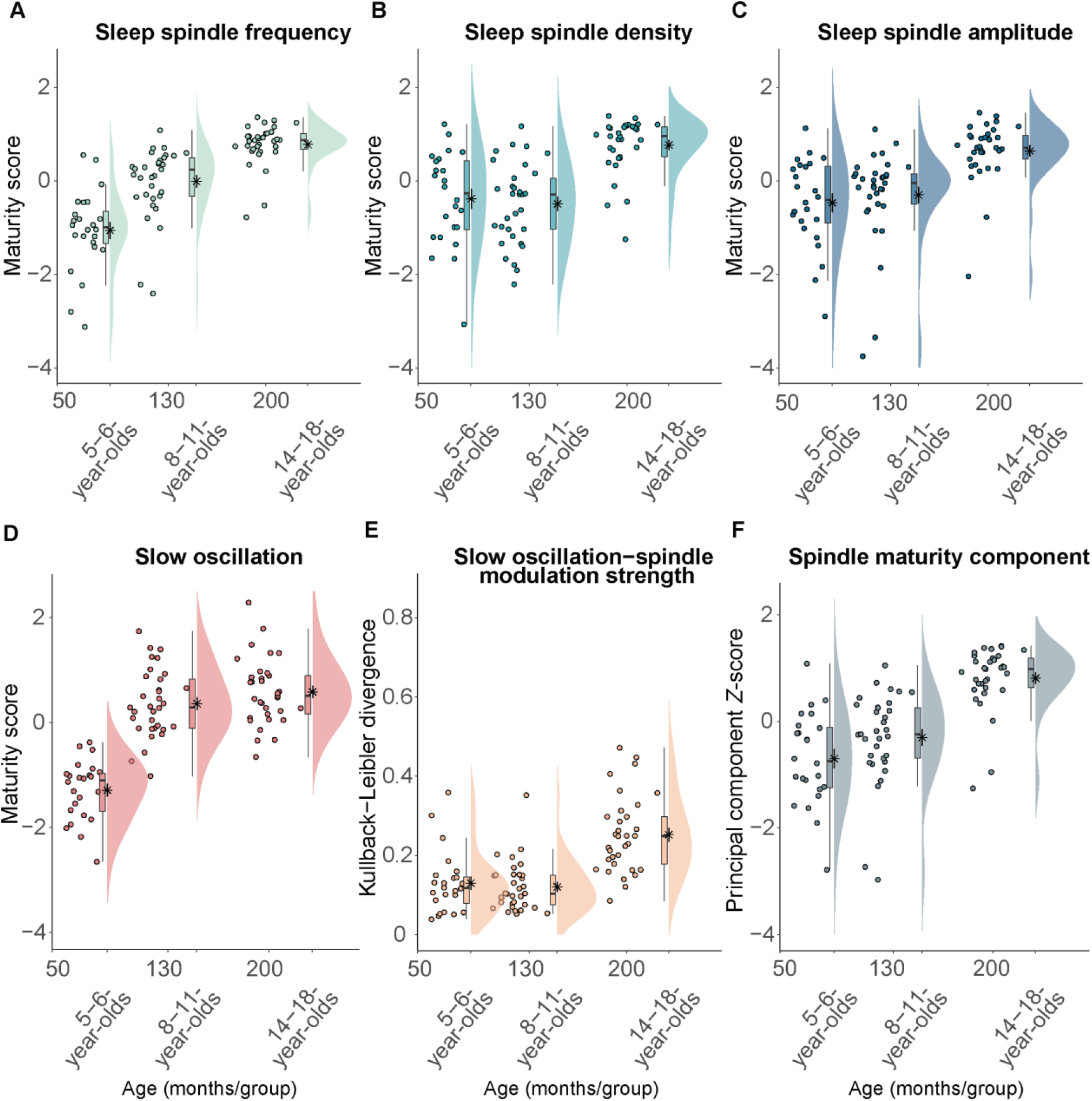
Measures of sleep spindle and slow oscillation maturity and slow oscillation-spindle coupling strength across age groups. (A-C) Fast sleep spindle maturity scores for (A) frequency, (B) density, and (C) amplitude. Maturity scores reflect the *Z*-standardized differences between adult-like and development-specific fast centro-parietal SPs. (D) Slow oscillation maturity scores represent the *Z*-standardized difference between frontal and centro-parietal amplitudes. (E) Kullback-Leibler divergence for development-specific fast centro-parietal sleep spindle modulation during centro-parietal slow oscillations, reflecting slow oscillation-spindle modulation strength. (F) First principal component of a principal component analysis onthethreefast spindlematurity scores which are shown in (A-C). Forall measures, highervalues are linked to olderage. Asterisks illustrate the mean.

Following a similar logic, we also created a measure for SO maturity. Based on our observation that age differences are mostly reflected in the emerging frontal dominance of SOs with older age (Figure 2) and on the literature (Kurth, Ringli, et al., 2010; Timofeev et al., 2020), we calculated the difference between the amplitude in frontal and centro-parietal regions to reflect the maturity pattern of SOs. In parallel with the procedure for the fast SP maturity scores, we *Z*-standardized the SO difference measure to be in the same metric space as the SP maturity values for all subsequent analyses. Comparably to the fast SP maturity scores, participants of older age showed higher *Z*-standardized SO difference scores (Figure 5D).

To examine how fast SP maturity would be associated with SO-SP coupling, we conducted two analyses. For one, we examined the relation between the pattern of power modulations in the SP frequency range (9-16 Hz) during the complete SO trial (down peak ± 1.2 sec, cf. time-frequency *t*-maps, Figure 3) with the fast SP maturity scores for SP frequency, density, and amplitude using PLSC. This analysis provides SO-SP coupling profiles associated with specific multivariate patterns of SP maturity. Based on a permutation test, we identified one significant pair of SP maturity and SO-SP coupling patterns (singular vector pair, *p* < .001) reflecting one common correlation between all fast SP maturity measures (as indicated by the SP maturity profile, Figure 6A) and a specific time-frequency SO-SP coupling pattern (Figure 6B). As can be inferred from Figure 6B, the SO-SP coupling pattern suggests, that higher fast SP maturity (higher values in Figure 6A) was associated with a more adult-like SO-SP coupling pattern, reflected in: (i) lower canonical fast SP power and higher power in canonical slow SP range during the down state and (ii) and higher power in the canonical fast SP frequencies range during the SO up peak. Overall this indicates that a stronger presence of SPs with more adult-like fast SP characteristics is associated with the well-known pattern of increased canonical fast SP activity during the SO up peak, increased canonical slow SP activity during the down state, and decreased activity of canonical fast SPs during the down peak.

**Figure 6.**
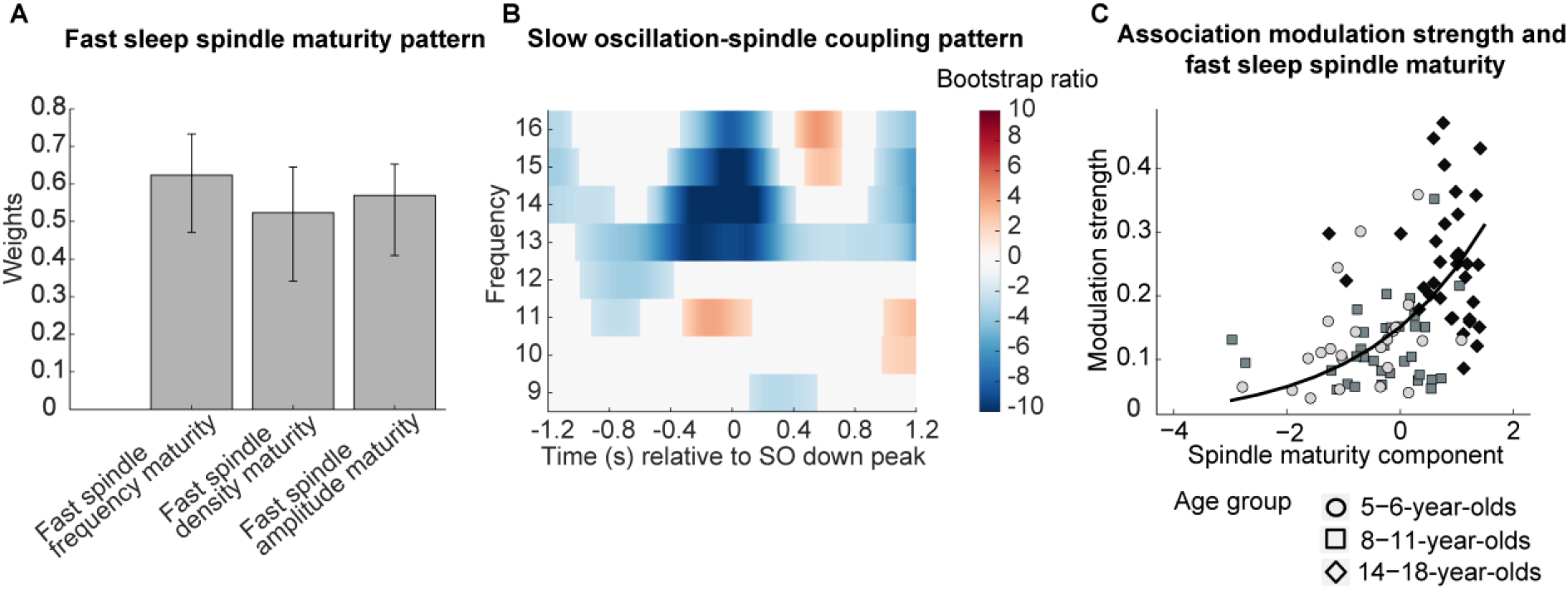
Association between fast spindle maturity and centro-parietal slow oscillation-development-specific fast spindle coupling. (A-B) Results from a partial least squares correlation revealed one (A) fast spindle maturity profile significantlyassociated with (B) a centro-parietal slow oscillation-spindle coupling pattern. (A) Weights of the first singular vector dimension of the fast spindle maturity scores. Error bars represent 95% bootstrap confidence intervals. (B) Weights of the first singular vector dimension of the slow oscillation-spindle coupling pattern by means of bootstrap ratios. Only values >1.96 and <-1.96 are colored. (C) Scatterplot of the association between the modulation strength (KL divergence) of development-specific fast centro-parietal SPs during centro-parietal SOs and the fast spindle maturity component. The curved line represents the prediction from the generalized linear mixed-effects model for the simple effect of the spindle maturity component. For visualization purposes, thethree agegroups are indicated bydifferent shapes and colors. Results for frontal slow oscillations can be found in Supplementary Figure 7.

After having identified that fast SP maturity was associated with more precise modulation of SP power during the SO cycle, we further examined whether fast SP maturity would also be related to the strength of modulation of development-specific fast centro-parietal SPs during SOs. Since all three indicators of fast SP maturity (frequency, density, amplitude maturity) showed age differences and were related to the pattern of SP power modulations during SOs, we used PCA to create one latent component capturing the maximal amount of variance across our fast SP maturity indicators (as for individual SP maturity scores, higher component values indicate higher fast SP maturity, Figure 5F). This fast SP maturity component was then used to examine the relation between fast SP maturity and development-specific fast SP modulation strength. We quantified the strength of modulation of development-specific fast centro-parietal SPs during centro-parietal SOs using the KL divergence (see Figure 5E, higher values are associated with older age). We then conducted a log-linked gamma GLMM analysis with the KL divergence during centro-parietal SOs as the dependent variable and the fast SP maturity component as a fixed effect, allowing for a random intercept for each participant. We further added the SO maturity score as a covariate. The GLMM revealed that fast SP maturity was uniquely associated with stronger modulation of development-specific fast centro-parietal SPs during SOs (β = - 0.49, *t* = −7.44, *p* < .001; see Figure 6C), while SO maturity was not significantly related to the modulation strength (β = 0.08, *t* = 1.04, *p* = .300). In sum, our results suggest that developmental age differences in the expression of SO-SP coupling depend on the degree to which thedominant fast SP typewithin an individual, i.e., the SPs that can be identified as peaks in the power spectrum (Figure 1A; Aru et al., 2015), shares characteristics with canonical fast SPs.

## 4 Discussion

Although the synchronization of SPs by the up peak of SOs and its functional significance has been recognized for decades, its development remains elusive (Muehlroth et al., 2019; Schreiner et al., 2021; Staresina et al., 2015; Steriade, 2006). Based on within-person detection of single events, we provide a detailed characterization of age-specific patterns of slow and fast SPs, SOs, and their coupling in children aged 5 to 6, 8 to 11, and 14 to 18 years. Specifically, in the youngest age group, we noted that the predominant type of fast SPs, as characterized by peaks in the power spectrum, is found in a frequency range slower than known from research in adults. Surprisingly, the inspection of SO-SP coupling patterns suggested synchronization being driven by SPs in the canonical fast SP range – even in the younger age groups but notably less precisely. Additional single event detection restricted to the canonical fast SP range indeed revealed adult-like fast centro-parietal SP events in all age groups, although with only minor presence in the child cohorts. Interrogation of coupling precision by means of PETHs confirmed that the coupling pattern found in adults, i.e., reduced SP occurrence during the down peak and increased SP likelihood precisely during the up state, can only be found for adult-like fast SPs and is less pronounced for the prevailing development-specific SPs found in the younger age groups.

To further corroborate these observations, we determined a personalized measure for fast SP maturity based on the differences between the predominant development-specific and the adult-like fast SPs in terms of frequency, amplitude, and density. Indeed, we observed that fast SP maturity (i.e., higher SP maturity scores) was associated with a more precise SO-SP modulation pattern of (i) decreased canonical fast SP power during the down state, (ii) increased canonical fast SP power during the up state, and (iii) increased canonical slow SP power in the transition from the up to the down peak. Most importantly, more advanced fast SP maturity exclusively was linked to stronger modulation of development-specific fast centro-parietal SPs during SOs, over and above SO maturity. Hence, the present results provide evidence that the development of precise SO-SP coupling may specifically be linked to the maturation of fast SPs.

Overall, our findings suggest that the precise coupling of SPs to the up peak of SOs might not be an inherent feature of the thalamocortical system. Rather, SO-SP coupling differs systematically across the lifespan (Hahn et al., 2020; Helfrich et al., 2018; Joechner et al., 2021; Muehlroth et al., 2019), likely depending on age-specific anatomical and electrophysiological properties of the thalamocortical network.

Previous research in older adults suggested that age-related dispersion and imprecision of frontal SO-SP coupling were related to atrophy in the prefrontal cortex and the thalamus – brain sites critically involved in the generation of mature SOs and SPs, respectively (Helfrich et al., 2018; Muehlroth et al., 2019). While these studies indirectly hypothesized that age-related changes in fast SP and SO features may contribute to alterations in the SO-SP coupling pattern, we provide direct evidence that the development of the well-known coupling of SPs to SO up peaks is uniquely linked to the emergence of an increasing number of SPs in the canonical fast SP frequency range. However, the exact anatomical and functional developments underlying the emergence of canonical fast SPs and their modulation by SOs remain to be elucidated. Furthermore, the question arises whether development-specific and adult-like fast SPs represent distinct representations of the same though developing generating network or whether they originate through different structures and connections.

SPs arise within well-described thalamic nuclei and thalamocortical circuits (Fernandez & Lüthi, 2020; Steriade, 1995). For one, the duration of hyperpolarization and the resulting length of hyperpolarization-rebound sequences in thalamocortical cells has been reported to particularly account for slower and faster SP frequencies (Steriade, 2003; Steriade & Llinás, 1988). Hence, decreasing hyperpolarization in thalamocortical, and potentially reticular thalamic and cortical, cells may underly the global increase in the frequency of predominant SPs across child and adolescent development (Campbell & Feinberg, 2016; Zhang et al., 2021). A relatively stronger age-related drop in hyperpolarization in topographically selected assemblies may lead to the expression of canonical fast SPs over posterior scalp electrodes. While the overall state of the brain determines the excitability of thalamocortical cells (Steriade & Llinás, 1988), it is still elusive which cellular or other developments could drive age-related alterations in thalamocortical hyperpolarizations.

Further, both thalamic nuclei and thalamocortical connectivity are refined across development (Steiner et al., 2020). This applies to structural and functional changes with studies indicating increased white matter and functional connectivity of thalamocortical tracts with older age – specifically with frontal, motor, and somatosensory cortical areas (Avery et al., 2021; Fair et al., 2010; Steiner et al., 2020). Previously, inter-individual differences in magnetic resonance 25 imaging indicators of white matter properties, supposed to index myelin-dependent efficiency in signal transmission (Chanraud et al., 2010), were linked to the expression of SP power and frequency (Mander et al., 2017; Piantoni et al., 2013; Sanchez et al., 2020; Vien et al., 2019). Therefore, increased white matter integrity of certain thalamocortical projections may support the increase in SP frequency and higher numbers of fast SPs with older age during development.

Importantly, accumulating findings support the conjecture that primarily canonical fast SPs coalesce with the up peak of SOs, while slow SPs are synchronized more towards the down peak (Bastian et al., 2022; Kurz et al., 2021; Mölle et al., 2011; Muehlroth et al., 2019). If merely the absolute frequency range of SPs would determine the coupling pattern, one might have simply expected to find a phase shift in coupling with older age and accompanying faster SPs. However, in neither age group, we did observe strong evidence indicating that SPs outside of the canonical fast SP frequency range or at other topographic locations couple to distinct phases of the SO. Rather than a synchronization of development-specific fast SPs during the down state, we found a general pattern of lower SP modulation in younger children where development-specific fast SP frequency is the lowest. However, development-specific fast SPs were not only lower in frequency but also occurred less often in younger children. Surprisingly though, the modulation of adult-like fast SPs during SOs was well detectable across all age groups with this co-occurrence pattern becoming more precise with older age – despite huge age-related differences in their occurrence rate and the fact that adult-like fast SPs showed the lowest density in all child cohorts (see Figure 1). Therefore, our data suggest that neither frequency nor density alone explains the SO-SP coupling pattern. Rather, in individuals increasingly capable of expressing adult-like fast SPs, SPs at all frequencies are globally modulated more strongly and in a comparable fashion. However, it remains an open question how the distinct coalescence of canonical fast and slow SPs with SOs develops.

While SPs can be solely initiated in intra-thalamic circuits and thus could co-occur with SOs merely by chance, corticothalamic input is one of the most potent mechanisms inducing SPs (Bonjean et al., 2011; Contreras & Steriade, 1995; Helfrich et al., 2018). The connectivity between the cortex and the thalamus is supposed to explain the formation of SO-SP modulation in a way that the synchronous firing of cortical cells during the depolarizing SO up state can excite thalamic SP generation while the joint neuronal hyperpolarization during the down state terminates and inhibits SPs (Contreras & Steriade, 1995; Steriade, 2006). Hence, it is conceivable that enhanced SO-SP coupling not only indexes the development of thalamocortical pathways (that may give rise to SPs at faster frequencies) but also changes in corticothalamic connectivity allowing for increasingly efficient cortical control over SP generation – specifically of canonical fast SPs (Contreras & Steriade, 1995).

While in our analyses SO maturity did not significantly explain SO-SP coupling development over and above fast SP maturity, there is evidence indicating that SO characteristics are associated with the pattern of SO-SP co-occurrence. For instance, a higher SO amplitude was associated with coupling strength (Kurz et al., 2021). Further, in older adults, a diminished directional influence of SOs was related to imprecise SO-SP coupling (Helfrich et al., 2018). Therefore, we cannot exclude a potentially important contribution of the development and characteristics of SOs for SO-SP coupling. It may just be that the prominent changes related to fast SPs across childhood and adolescence may be relatively more influential and/or that SO features other than the ones examined here may be effective in shaping SO-SP coupling.

### 4.1 Conclusion

Overall, our findings describe a unique relation between fast SP maturation and the development of SO-SP coupling. While SOs provide optimal time windows for SPs to arise, it is the ability to generate adult-like fast SPs that determines coupling strength and precision across child and adolescent development. Given the evidence for its generating role, our results implicate the maturation of specific thalamocortical circuits as the cornerstone of adult-like SO-SP coupling patterns. Hence, our findings represent a promising starting point for future research addressing the precise relation between age-related changes in brain structure and function, the emergence of adult-like SO-SP coupling, and cognitive development across childhood and adolescence.

## Funding statement and acknowledgments

This research was conducted within the project “Lifespan Rhythms of Memory and Cognition” (RHYME, PI: MW-B) at the Max Planck Institute for Human Development (MPIB), Berlin, Germany, and at the Department of Psychology, Laboratory for Sleep, Cognition and Consciousness Research, University of Salzburg, Salzburg, Austria (PI: KH). A-KJ is a fellow of the International Max Planck Research School on the Life Course (LIFE; http://www.imprs-life.mpg.de/en). MW-B received support from the German Research Foundation (DFG, WE 4269/5-1) and the Jacobs Foundation (Early Career Research Fellowship 2017–2019). KH was supported by Austrian Science Fund (T397-B02, P25000), the Jacobs Foundation (JS1112H), and the Centre for Cognitive Neuroscience Salzburg (CCNS). MAH was additionally supported by the Doctoral College “Imaging the Mind” (FWF, Austrian Science Fund W1233-G17). We thank S. Wehmeier for her invaluable help with data collection at the MPIB. We are grateful to all members of the RHYME and LIME projects for valuable feedback. We further acknowledge support by the Max Planck Dahlem Campus of Cognition (MPDCC). Finally, we thank all our participants and their families for their time as well as the principals of the schools and the local education authority (Mag. Dipl. Paed. B. Heinrich, Prof. Mag. J. Thurner) in Salzburg who supported this research.

