## Supplemental Information for "Sleep spindle maturation enhances slow oscillation-spindle coupling"

Supplementary Table 1

*Sleep architecture*

| | 5–6-year-olds | | | 8–11-year-olds | | | 14–18-year-olds | | |
|---|---|---|---|---|---|---|---|---|---|
| | Mean | Median | Percentile [25%; 75%] | Mean | Median | Percentile [25%; 75%] | Mean | Median | Percentile [25%; 75%] |
| TST (min) | 605.52 | 610.00 | [593.88; 629.12] | 560.44 | 565.50 | [540.50; 590.00] | 474.70 | 486.50 | [440.50; 505.00] |
| N1(%) | 5.95 | 5.19 | [3.96;7.33] | 2.49 | 2.21 | [1.32; 3.36] | 8.69 | 7.60 | [5.26; 11.17] |
| N2(%) | 35.88 | 35.40 | [34.06; 39.20] | 43.62 | 42.87 | [39.15; 49.78] | 40.70 | 39.85 | [37.63; 43.27] |
| N3(%) | 24.73 | 23.82 | [21.21; 26.96] | 26.61 | 26.46 | [21.78; 32.57] | 25.38 | 23.65 | [20.80; 28.09] |
| R (%) | 33.44 | 33.67 | [30.27; 34.80] | 27.27 | 25.86 | [22.01; 31.08] | 25.24 | 25.42 | [21.21; 29.64] |
| WASO (min) | 10.33 | 5.75 | [4.00; 9.12] | 10.06 | 6.50 | [2.50; 13.00] | 10.55 | 9.00 | [4.00; 16.00] |
| NREM (%) | 60.61 | 61.15 | [56.48; 64.50] | 70.23 | 71.48 | [67.03; 75.12] | 66.07 | 65.49 | [62.54; 70.60] |

*Note*. TST=Total sleep time, WASO=Wake after sleep onset, R=rapid-eye movement sleep, NREM=sum of N2 and N3 sleep.

Supplementary Table 2

*Type III analysis of variance table from a linear mixed-effects model on the effects of age group and slow oscillation topography on the co-occurrence of slow frontal sleep spindles with slow oscillations*

| Predictor | Sum of squares | Mean square | $df_{\text{Num}}$ | $df_{\text{Den}}$ | $F$ | $p$ |
|---|---|---|---|---|---|---|
| Age group | 121.10 | 60.55 | 2 | 86.32 | 16.66 | < .001 |
| Topography | 1,076.71 | 538.35 | 2 | 204.56 | 148.13 | < .001 |
| Age group*topography | 452.79 | 113.20 | 4 | 204.56 | 31.15 | < .001 |

*Note*. Model formula: measurement ~ age group * topography + (1 | ID), F_SPCoup_Data, TRUE, lmerControl(optimizer = "nloptwrap", optCtrl = list(algorithm = "NLOPT_LN_BOBYQA")), na.exclude. $df_{\text{Num}}$ indicates the degrees of freedom numerator. $df_{\text{Den}}$ indicates the degrees of freedom denominator. Degrees of freedom were determined using Satterthwaite's method.

Supplementary Table 3

*Type III analysis of variance table from a linear mixed-effects model on the effects of age group and slow oscillation topography on the co-occurrence of development-specific fast centro-parietal sleep spindles with slow oscillations*

| Predictor | Sum of squares | Mean square | $df_{Num}$ | $df_{Den}$ | F | p |
|---|---|---|---|---|---|---|
| Age group | 39.87 | 19.93 | 2 | 91.53 | 6.05 | .003 |
| Topography | 1,056.77 | 528.39 | 2 | 206.95 | 160.4 | < .001 |
| Age group*topography | 322.32 | 80.58 | 4 | 206.95 | 24.46 | < .001 |

*Note*. Model formula: measurement ~ age group * topography + (1 | ID), CP_SPCoup_Data, TRUE, lmerControl(optimizer = “nloptwrap”, optCtrl = list(algorithm = “NLOPT_LN_BOBYQA”)), na.exclude. $df_{Num}$ indicates the degrees of freedom numerator. $df_{Den}$ indicates the degrees of freedom denominator. Degrees of freedom were determined using Satterthwaite’s method.

Supplementary Table 4

*Type III analysis of variance table from a linear mixed-effects model on the effects of age group and slow oscillation topography on the co-occurrence of slow oscillations with slow frontal sleep spindles*

| Predictor | Sum of squares | Mean square | $df_{Num}$ | $df_{Den}$ | $F$ | $p$ |
|---|---|---|---|---|---|---|
| Age group | 1.04 | 0.52 | 2 | 87.31 | 0.75 | .476 |
| Topography | 69.63 | 34.82 | 2 | 206.09 | 49.95 | < .001 |
| Age group*topography | 20.08 | 5.02 | 4 | 206.09 | 7.20 | < .001 |

*Note*. Model formula: lmer, measurement ~ age group * topography + (1 | ID), SOCoup_Data_FSP, TRUE, lmerControl(optimizer = “nloptwrap”, optCtrl = list(algorithm = “NLOPT_LN_BOBYQA”)), na.exclude. $df_{Num}$ indicates the degrees of freedom numerator. $df_{Den}$ indicates the degrees of freedom denominator. Degrees of freedom were determined using Satterthwaite’s method.

Supplementary Table 5

*Type III analysis of variance table from a linear mixed-effects model on the effects of age group and slow oscillation topography on the co-occurrence of slow oscillations with development-specific fast centro-parietal sleep spindles*

| Predictor | Sum of squares | Mean square | $df_{Num}$ | $df_{Den}$ | $F$ | $p$ |
|---|---|---|---|---|---|---|
| Age group | 6.20 | 3.10 | 2 | 91.05 | 4.88 | .010 |
| Topography | 83.92 | 41.96 | 2 | 208.02 | 66.12 | < .001 |
| Age group*topography | 15.67 | 3.92 | 4 | 208.02 | 6.17 | < .001 |

*Note*. Model formula: lmer, measurement ~ age group * topography + (1 | ID), SOCoup_Data_CPSP, TRUE, lmerControl(optimizer = “nloptwrap”, optCtrl = list(algorithm = “NLOPT_LN_BOBYQA”)), na.exclude. $df_{Num}$ indicates the degrees of freedom numerator. $df_{Den}$ indicates the degrees of freedom denominator. Degrees of freedom were determined using Satterthwaite’s method.

Supplementary Table 6

*Type III analysis of variance table from a linear mixed-effects model on the effects of age group and slow oscillation topography on the co-occurrence of adult-like fast centro-parietal sleep spindles with slow oscillations*

| Predictor | Sum of squares | Mean square | $df_{Num}$ | $df_{Den}$ | $F$ | $p$ |
|---|---|---|---|---|---|---|
| Age group | 256.17 | 128.09 | 2 | 89.56 | 20.29 | < .001 |
| Topography | 1,885.45 | 942.73 | 2 | 206.11 | 149.3 | < .001 |
| Age group*topography | 481.67 | 120.42 | 4 | 206.11 | 19.08 | < .001 |

*Note*. Model formula: measurement ~ age group * topography + (1 | ID), SPCoup_Data_adult, TRUE, lmerControl(optimizer = "nloptwrap", optCtrl = list(algorithm = "NLOPT_LN_BOBYQA")), na.exclude. $df_{Num}$ indicates the degrees of freedom numerator. $df_{Den}$ indicates the degrees of freedom denominator. Degrees of freedom were determined using Satterthwaite's method.

Supplementary Table 7

*Type III analysis of variance table from a linear mixed-effects model on the effects of age group and slow oscillation topography on the co-occurrence of slow oscillations with adult-like fast centro-parietal sleep spindles*

| Predictor | Sum of squares | Mean square | $df_{Num}$ | $df_{Den}$ | $F$ | $p$ |
|---|---|---|---|---|---|---|
| Age group | 19.73 | 9.86 | 2 | 91.70 | 14.78 | < .001 |
| Topography | 80.13 | 40.07 | 2 | 208.39 | 60.05 | < .001 |
| Age group*topography | 15.43 | 3.86 | 4 | 208.39 | 5.78 | < .001 |

*Note*. Model formula: measurement ~ age group * topography + (1 | ID), SOCoup_Data_adult, TRUE, lmerControl(optimizer = "nloptwrap", optCtrl = list(algorithm = "NLOPT_LN_BOBYQA")), na.exclude. *df*=degrees of freedom. $df_{Num}$ indicates the degrees of freedom numerator. $df_{Den}$ indicates the degrees of freedom denominator. Degrees of freedom were determined using Satterthwaite's method.

Supplementary Table 8

*Type III analysis of variance table from a linear mixed-effects model on the effects of age group and sleep spindle topography on sleep spindle spectral peak frequency*

| Predictor | Sum of squares | Mean square | $df_{Num}$ | $df_{Den}$ | $F$ | $p$ |
|---|---|---|---|---|---|---|
| Age group | 17.26 | 8.63 | 2 | 79.77 | 69.64 | < .001 |
| Topography | 8.56 | 8.56 | 1 | 120.05 | 69.13 | < .001 |
| Age group*topography | 0.17 | 0.08 | 2 | 120.05 | 0.68 | .507 |

*Note*. $df_{Num}$ indicates the degrees of freedom numerator. $df_{Den}$ indicates the degrees of freedom denominator. Degrees of freedom were determined using Satterthwaite's method.

Supplementary Table 9

*(A)Post-hoc comparisons for the topography effect on sleep spindle peak frequency based on estimated marginal means (B) Post-hoc comparisons for the age group effect on sleep spindle peak frequency based on estimated marginal means*

A

| Contrast | Estimate | *SE* | *df* | *t* | $p_{\text{adj}}$ |
|---|---|---|---|---|---|
| Frontal – Centro-parietal | -0.44 | 0.05 | 120.05 | -8.31 | < .001 |

B

| Contrast | Estimate | *SE* | *df* | *t* | $p_{adj}$ |
|---|---|---|---|---|---|
| 5–6-year-olds – 8–11-year-olds | -0.46 | 0.12 | 68.35 | -3.85 | .001 |
| 5–6-year-olds – 14–18-year-olds | -1.06 | 0.12 | 68.35 | -8.88 | < .001 |
| 8–11-year-olds – 14–18-year-olds | -0.60 | 0.06 | 120.05 | -9.79 | < .001 |

*Note*. Degrees of freedom were determined using Satterthwaite's method. $P_{adj}$-values are Bonferroni corrected. *SE*= standard error, *df*= degrees of freedom.

Supplementary Table 10

*Type III analysis of variance table from a linear mixed-effects model on the effects of age group and sleep spindle topography on sleep spindle frequency*

| Predictor | Sum of squares | Mean square | $df_{\text{Num}}$ | $df_{\text{Den}}$ | *F* | *p* |
|---|---|---|---|---|---|---|
| Age group | 19.34 | 9.67 | 2 | 77.74 | 113.76 | < .001 |
| Topography | 11.93 | 11.93 | 1 | 119.51 | 140.39 | < .001 |
| Age group*topography | 0.15 | 0.07 | 2 | 119.51 | 0.86 | .425 |

*Note*. $df_{\text{Num}}$ indicates the degrees of freedom numerator. $df_{\text{Den}}$ indicates the degrees of freedom denominator. Degrees of freedom were determined using Satterthwaite's method.

Supplementary Table 11

*(A)Post-hoc comparisons for the topography effect on sleep spindle frequency based on estimated marginal means (B) Post-hoc comparisons for the age group effect on sleep spindle frequency based on estimated marginal means*

| A | | | | | |
|---|---|---|---|---|---|
| Contrast | Estimate | *SE* | *df* | *t* | $p_{adj}$ |
| Frontal – Centro-parietal | -0.52 | 0.04 | 119.51 | -11.85 | < .001 |
| **B** | | | | | |
| Contrast | Estimate | *SE* | *df* | *t* | $p_{adj}$ |
| 5–6-year-olds – 8–11-year-olds | -0.51 | 0.12 | 63.27 | -4.22 | < .001 |
| 5–6-year-olds – 14–18-year-olds | -1.18 | 0.12 | 63.27 | -9.83 | < .001 |
| 8–11-year-olds – 14–18-year-olds | -0.67 | 0.05 | 119.51 | -13.26 | < .001 |

*Note*. Degrees of freedom were determined using Satterthwaite's method. $P_{adj}$-values are Bonferroni corrected. *SE*= standard error, *df*= degrees of freedom.

Supplementary Table 12

*Type III analysis of variance table from a linear mixed-effects model on the effects of age group and sleep spindle topography on sleep spindle density*

| Predictor | Sum of squares | Mean square | $df_{Num}$ | $df_{Den}$ | *F* | *p* |
|---|---|---|---|---|---|---|
| Age group | 1.43 | 0.71 | 2 | 77.27 | 14.47 | < .001 |
| Topography | 2.04 | 2.04 | 1 | 119.22 | 41.26 | < .001 |
| Age group*topography | 0.73 | 0.36 | 2 | 119.22 | 7.35 | .001 |

*Note*. $df_{Num}$ indicates the degrees of freedom numerator. $df_{Den}$ indicates the degrees of freedom denominator. Degrees of freedom were determined using Satterthwaite's method.

Supplementary Table 13

*Post-hoc comparisons for the sleep spindle density interaction effect based on estimated marginal means*

| Contrast | Estimate | *SE* | *df* | *t* | $p_{adj}$ |
|---|---|---|---|---|---|
| 5–6-y F – 8–11-y F | -0.43 | 0.10 | 87.16 | -4.13 | **.001** |
| 5–6-y F – 14–18-y F | -0.22 | 0.10 | 87.16 | -2.14 | .529 |
| 5–6-y F – 5–6-y CP | 0.36 | 0.06 | 119.22 | 5.54 | **< .001** |
| 5–6-y F – 8–11-y CP | -0.18 | 0.10 | 87.16 | -1.74 | 1.00 |
| 5–6-y F – 14–18-y CP | -0.18 | 0.10 | 87.16 | -1.71 | 1.00 |
| 8–11-y F – 14–18-y F | 0.21 | 0.05 | 119.22 | 3.75 | **.004** |
| 8–11-y F – 5–6-y CP | 0.78 | 0.10 | 87.16 | 7.58 | **< .001** |
| 8–11-y F – 8–11-y CP | 0.25 | 0.05 | 119.22 | 4.50 | **< .001** |
| 8–11-y F – 14–18-y CP | 0.25 | 0.05 | 119.22 | 4.56 | **< .001** |
| 14–18-y F – 5–6-y CP | 0.58 | 0.10 | 87.16 | 5.59 | **< .001** |
| 14–18-y F – 8–11-y CP | 0.04 | 0.05 | 119.22 | 0.75 | 1.00 |
| 14–18-y F – 14–18-y CP | 0.04 | 0.05 | 119.22 | 0.81 | 1.00 |
| 5–6-y CP – 8–11-y CP | -0.53 | 0.10 | 87.16 | -5.19 | **< .001** |
| 5–6-y CP –14–18-y CP | -0.53 | 0.10 | 87.16 | -5.16 | **< .001** |
| 8–11-y CP – 14–18-y CP | 0.00 | 0.05 | 119.22 | 0.06 | 1.00 |

*Note*. Degrees of freedom were determined using Satterthwaite's method. $P_{adj}$-values are Bonferroni corrected. *SE*= standard error, *df*= degrees of freedom. F=frontal, CP= centro-parietal.

Supplementary Table 14

*Type III analysis of variance table from a linear mixed-effects model on the effects of age group and sleep spindle topography on sleep spindle amplitude*

| Predictor | Sum of squares | Mean square | $df_{Num}$ | $df_{Den}$ | *F* | *p* |
|---|---|---|---|---|---|---|
| Age group | 3,494.22 | 1,747.11 | 2 | 79.62 | 48.51 | < .001 |
| Topography | 8,103.15 | 8,103.15 | 1 | 120.25 | 225.00 | < .001 |
| Age group*topography | 363.57 | 181.78 | 2 | 120.25 | 5.05 | .008 |

*Note*. $df_{Num}$ indicates the degrees of freedom numerator. $df_{Den}$ indicates the degrees of freedom denominator. Degrees of freedom were determined using Satterthwaite's method.

Supplementary Table 15

*Post-hoc comparisons for the sleep spindle amplitude interaction effect based on estimated marginal means*

| Contrast | Estimate | *SE* | *df* | *t* | $p_{adj}$ |
|---|---|---|---|---|---|
| 5–6-y F – 8–11-y F | -8.18 | 2.41 | 103.18 | -3.39 | **.015** |
| 5–6-y F – 14–18-y F | 5.04 | 2.41 | 103.18 | 2.09 | .586 |
| 5–6-y F – 5–6-y CP | 11.54 | 1.73 | 120.25 | 6.66 | **< .001** |
| 5–6-y F – 8–11-y CP | 9.33 | 2.41 | 103.18 | 3.87 | **.003** |
| 5–6-y F – 14–18-y CP | 16.70 | 2.41 | 103.18 | 6.93 | **< .001** |
| 8–11-y F – 14–18-y F | 13.21 | 1.48 | 120.25 | 8.94 | **< .001** |
| 8–11-y F – 5–6-y CP | 19.72 | 2.41 | 103.18 | 8.18 | **< .001** |
| 8–11-y F – 8–11-y CP | 17.51 | 1.48 | 120.25 | 11.85 | **< .001** |
| 8–11-y F – 14–18-y CP | 24.87 | 1.48 | 120.25 | 16.84 | **< .001** |
| 14–18-y F – 5–6-y CP | 6.51 | 2.41 | 103.18 | 2.70 | .121 |
| 14–18-y F – 8–11-y CP | 4.29 | 1.48 | 120.25 | 2.91 | .065 |
| 14–18-y F – 14–18-y CP | 11.66 | 1.48 | 120.25 | 7.89 | **< .001** |
| 5–6-y CP – 8–11-y CP | -2.21 | 2.41 | 103.18 | -0.92 | 1.00 |
| 5–6-y CP – 14–18-y CP | 5.15 | 2.41 | 103.18 | 2.14 | .524 |
| 8–11-y CP – 14–18-y CP | 7.37 | 1.48 | 120.25 | 4.99 | **< .001** |

*Note*. Degrees of freedom were determined using Satterthwaite's method. $P_{adj}$-values are Bonferroni corrected. *SE*= standard error, *df*= degrees of freedom. F=frontal, CP= centro-parietal.

Supplementary Table 16

*Type III analysis of variance table from a linear mixed-effects model on the effects of age group and sleep spindle type (development-specific and adult-like fast sleep spindles) on frequency*

| Predictor | Sum of squares | Mean square | $df_{\text{Num}}$ | $df_{\text{Den}}$ | *F* | *p* |
|---|---|---|---|---|---|---|
| Age group | 9.76 | 4.88 | 2 | 87.41 | 55.91 | < .001 |
| Spindle type | 38.70 | 38.70 | 1 | 124.05 | 443.39 | < .001 |
| Age group*spindle type | 9.53 | 4.76 | 2 | 124.05 | 54.58 | < .001 |

*Note*. $df_{\text{Num}}$ indicates the degrees of freedom numerator. $df_{\text{Den}}$ indicates the degrees of freedom denominator. Degrees of freedom were determined using Satterthwaite's method.

Supplementary Table 17

*Post-hoc comparisons for development-specific and adult-like fast sleep spindle frequency values within the three age groups (interaction effect) based on estimated marginal means*

| Contrast | Estimate | *SE* | *df* | *t* | $p_{\text{adj}}$ |
|---|---|---|---|---|---|
| 5–6-y adult-like – 8–11-y adult-like | 0.21 | 0.10 | 137.51 | 2.15 | .502 |
| 5–6-y adult-like – 14–18-y adult-like | -0.02 | 0.10 | 137.51 | -0.23 | 1.00 |
| 5–6-y adult-like – 5–6-y development-specific | 1.55 | 0.09 | 124.05 | 18.19 | **< .001** |
| 5–6-y adult-like – 8–11-y development-specific | 1.09 | 0.10 | 137.51 | 11.10 | **< .001** |
| 5–6-y adult-like – 14–18-y development-specific | 0.36 | 0.10 | 137.51 | 3.62 | **.006** |
| 8–11-y adult-like – 14–18-y adult-like | -0.23 | 0.07 | 124.05 | -3.23 | **.024** |
| 8–11-y adult-like – 5–6-y development-specific | 1.34 | 0.10 | 137.51 | 13.58 | **< .001** |
| 8–11-y adult-like – 8–11-y development-specific | 0.88 | 0.07 | 124.05 | 12.13 | **< .001** |
| 8–11-y adult-like – 14–18-y development-specific | 0.15 | 0.07 | 124.05 | 2.00 | .719 |
| 14–18-y adult-like – 5–6-y development-specific | 1.57 | 0.10 | 137.51 | 15.96 | **< .001** |
| 14–18-y adult-like – 8–11-y development-specific | 1.12 | 0.07 | 124.05 | 15.36 | **< .001** |

| Contrast | Estimate | *SE* | *df* | *t* | $p_{adj}$ |
|---|---|---|---|---|---|
| 14–18-y adult-like – 14–18-y development-specific | 0.38 | 0.07 | 124.05 | 5.22 | **< .001** |
| 5–6-y development-specific – 8–11-y development-specific | -0.46 | 0.10 | 137.51 | -4.63 | **< .001** |
| 5–6-y development-specific – 14–18-y development-specific | -1.19 | 0.10 | 137.51 | -12.11 | **< .001** |
| 8–11-y development-specific – 14–18-y development-specific | -0.74 | 0.07 | 124.05 | -10.14 | **< .001** |

*Note*. Degrees of freedom were determined using Satterthwaite's method. $P_{adj}$-values are Bonferroni corrected. *SE*= standard error, *df*= degrees of freedom.

Supplementary Table 18

*Type III analysis of variance table from a linear mixed-effects model on the effects of age group and sleep spindle type (development-specific and adult-like fast sleep spindles) on density*

| Predictor | Sum of squares | Mean square | $df_{Num}$ | $df_{Den}$ | *F* | *p* |
|---|---|---|---|---|---|---|
| Age group | 4.22 | 2.11 | 2 | 81.72 | 40.23 | < .001 |
| Spindle type | 5.62 | 5.62 | 1 | 122.60 | 107.28 | < .001 |
| Age group*spindle type | 1.85 | 0.92 | 2 | 122.60 | 17.65 | < .001 |

*Note*. $df_{Num}$ indicates the degrees of freedom numerator. $df_{Den}$ indicates the degrees of freedom denominator. Degrees of freedom were determined using Satterthwaite's method.

Supplementary Table 19

*Post-hoc comparisons for development-specific and adult-like fast sleep spindle density within the three age groups (interaction effect) based on estimated marginal means*

| Contrast | Estimate | *SE* | *df* | *t* | $p_{adj}$ |
|---|---|---|---|---|---|
| 5–6-y adult-like – 8–11-y adult-like | -0.50 | 0.10 | 97.35 | -5.02 | **< .001** |
| 5–6-y adult-like – 14–18-y adult-like | -0.93 | 0.10 | 97.35 | -9.38 | **< .001** |
| 5–6-y adult-like – 5–6-y development-specific | -0.48 | 0.07 | 122.60 | -7.23 | **< .001** |
| 5–6-y adult-like – 8–11-y development-specific | -1.01 | 0.10 | 97.35 | -10.21 | **< .001** |
| 5–6-y adult-like – 14–18-y development-specific | -1.01 | 0.10 | 97.35 | -10.18 | **< .001** |

| Contrast | Estimate | *SE* | *df* | *t* | $p_{adj}$ |
|---|---|---|---|---|---|
| 8–11-y adult-like – 14–18-y adult-like | -0.43 | 0.06 | 122.60 | -7.67 | **< .001** |
| 8–11-y adult-like – 5–6-y development-specific | 0.02 | 0.10 | 97.35 | 0.20 | 1.00 |
| 8–11-y adult-like – 8–11-y development-specific | -0.51 | 0.06 | 122.60 | -9.14 | **< .001** |
| 8–11-y adult-like – 14–18-y development-specific | -0.51 | 0.06 | 122.60 | -9.08 | **< .001** |
| 14–18-y adult-like – 5–6-y development-specific | 0.45 | 0.10 | 97.35 | 4.56 | **< .001** |
| 14–18-y adult-like – 8–11-y development-specific | -0.08 | 0.06 | 122.60 | -1.47 | 1.00 |
| 14–18-y adult-like – 14–18-y development-specific | -0.08 | 0.06 | 122.60 | -1.41 | 1.00 |
| 5–6-y development-specific – 8–11-y development-specific | -0.53 | 0.10 | 97.35 | -5.40 | **< .001** |
| 5–6-y development-specific – 14–18-y development-specific | -0.53 | 0.10 | 97.35 | -5.36 | **< .001** |
| 8–11-y development-specific – 14–18-y development-specific | 0.00 | 0.06 | 122.60 | 0.06 | 1.00 |

*Note*. Degrees of freedom were determined using Satterthwaite's method. $P_{adj}$-values are Bonferroni corrected. *SE*= standard error, *df*= degrees of freedom.

Supplementary Table 20

*Type III analysis of variance table from a linear mixed-effects model on the effects of age group and sleep spindle type (development-specific and adult-like fast sleep spindles) on amplitude*

| Predictor | Sum of squares | Mean square | $df_{Num}$ | $df_{Den}$ | *F* | *p* |
|---|---|---|---|---|---|---|
| Age group | 1,125.93 | 562.96 | 2 | 82.58 | 38.63 | < .001 |
| Spindle type | 419.32 | 419.32 | 1 | 122.53 | 28.77 | < .001 |
| Age group*spindle type | 114.91 | 57.45 | 2 | 122.53 | 3.94 | .022 |

*Note*. $df_{Num}$ indicates the degrees of freedom numerator. $df_{Den}$ indicates the degrees of freedom denominator. Degrees of freedom were determined using Satterthwaite's method.

Supplementary Table 21

*Post-hoc comparisons for development-specific and adult-like fast sleep spindle amplitude within the three age groups (interaction effect) based on estimated marginal means*

| Contrast | Estimate | *SE* | *df* | *t* | $p_{adj}$ |
|---|---|---|---|---|---|
| 5–6-y adult-like – 8–11-y adult-like | -2.76 | 1.50 | 108.89 | -1.84 | 1.00 |
| 5–6-y adult-like – 14–18-y adult-like | 1.55 | 1.50 | 108.89 | 1.03 | 1.00 |
| 5–6-y adult-like – 5–6-y development-specific | -4.47 | 1.10 | 122.53 | -4.06 | **.001** |
| 5–6-y adult-like – 8–11-y development-specific | -6.68 | 1.50 | 108.89 | -4.46 | **< .001** |
| 5–6-y adult-like – 14–18-y development-specific | 0.68 | 1.50 | 108.89 | 0.46 | 1.00 |
| 8–11-y adult-like – 14-18-y adult-like | 4.31 | 0.94 | 122.53 | 4.58 | **< .001** |
| 8–11-y adult-like – 5–6-y development-specific | -1.71 | 1.50 | 108.89 | -1.14 | 1.00 |
| 8–11-y adult-like – 8–11-y development-specific | -3.92 | 0.94 | 122.53 | -4.18 | **.001** |
| 8–11-y adult-like – 14–18-y development-specific | 3.44 | 0.94 | 122.53 | 3.66 | **.006** |
| 14–18-y adult-like – 5–6-y development-specific | -6.02 | 1.50 | 108.89 | -4.02 | **.002** |
| 14–18-y adult-like – 8–11-y development-specific | -8.23 | 0.94 | 122.53 | -8.76 | **< .001** |
| 14–18-y adult-like – 14–18-y development-specific | -0.87 | 0.94 | 122.53 | -0.92 | 1.00 |
| 5–6-y development-specific – 8–11-y development-specific | -2.21 | 1.50 | 108.89 | -1.48 | 1.00 |
| 5–6-y development-specific – 14–18-y development-specific | 5.15 | 1.50 | 108.89 | 3.44 | **.012** |
| 8–11-y development-specific – 14–18-y development-specific | 7.37 | 0.94 | 122.53 | 7.84 | **< .001** |

*Note*. Degrees of freedom were determined using Satterthwaite's method. $P_{adj}$-values are Bonferroni corrected. *SE*= standard error, *df*= degrees of freedom.

Supplementary Table 22

*Type III analysis of variance table from a linear mixed-effects model on the effects of age group and slow oscillation topography on slow oscillation frequency*

| Predictor | Sum of squares | Mean square | $df_{Num}$ | $df_{Den}$ | *F* | *p* |
|---|---|---|---|---|---|---|
| Age group | 0.03 | 0.01 | 2 | 94.24 | 65.87 | < .001 |
| Topography | 0.01 | 0.01 | 2 | 207.33 | 33.94 | < .001 |
| Age group*topography | 0.01 | 0.00 | 4 | 207.33 | 11.68 | < .001 |

*Note*. $df_{Num}$ indicates the degrees of freedom numerator. $df_{Den}$ indicates the degrees of freedom denominator. Degrees of freedom were determined using Satterthwaite's method.

Supplementary Table 23

*Post-hoc comparisons for the slow oscillation frequency interaction effect based on estimated marginal means*

| Contrast | Estimate | *SE* | *df* | *t* | $p_{adj}$ |
|---|---|---|---|---|---|
| 5–6-y F – 8–11-y F | -0.01 | 0.00 | 174.60 | -1.62 | 1.00 |
| 5–6-y F – 14–18-y F | 0.01 | 0.00 | 174.60 | 1.16 | 1.00 |
| 5–6-y F – 5–6-y CP | 0.00 | 0.00 | 207.33 | 0.14 | 1.00 |
| 5–6-y F – 8–11-y CP | -0.00 | 0.00 | 174.60 | -0.48 | 1.00 |
| 5–6-y F – 14–18-y CP | 0.01 | 0.00 | 174.60 | 2.89 | .158 |
| 5–6-y F – 5–6-y O | 0.01 | 0.00 | 207.33 | 1.39 | 1.00 |
| 5–6-y F – 8–11-y O | 0.00 | 0.00 | 174.60 | 0.43 | 1.00 |
| 5–6-y F – 14–18-y O | 0.04 | 0.00 | 174.60 | 8.72 | **< .001** |
| 8–11-y F – 14–18-y F | 0.01 | 0.00 | 207.33 | 3.78 | **.008** |
| 8–11-y F – 5–6-y CP | 0.01 | 0.00 | 174.60 | 1.74 | 1.00 |
| 8–11-y F – 8–11-y CP | 0.01 | 0.00 | 207.33 | 1.54 | 1.00 |
| 8–11-y F – 14–18-y CP | 0.02 | 0.00 | 207.33 | 6.11 | **< .001** |
| 8–11-y F – 5–6-y O | 0.01 | 0.00 | 174.60 | 2.82 | .194 |
| 8–11-y F – 8–11-y O | 0.01 | 0.00 | 207.33 | 2.78 | .215 |
| 8–11-y F – 14–18-y O | 0.05 | 0.00 | 207.33 | 14.02 | **< .001** |
| 14–18-y F – 5–6-y CP | -0.01 | 0.00 | 174.60 | -1.04 | 1.00 |
| 14–18-y F – 8–11-y CP | -0.01 | 0.00 | 207.33 | -2.24 | .950 |
| 14–18-y F – 14–18-y CP | 0.01 | 0.00 | 207.33 | 2.34 | .736 |
| 14–18-y F – 5–6-y O | 0.00 | 0.00 | 174.60 | 0.04 | 1.00 |
| 14–18-y F – 8–11-y O | -0.00 | 0.00 | 207.33 | -1.00 | 1.00 |
| 14–18-y F – 14–18-y O | 0.04 | 0.00 | 207.33 | 10.25 | **< .001** |
| 5–6-y CP – 8–11-y CP | -0.00 | 0.00 | 174.60 | -0.61 | 1.00 |
| 5–6-y CP – 14–18-y CP | 0.01 | 0.00 | 174.60 | 2.76 | .229 |
| 5–6-y CP – 5–6-y O | 0.01 | 0.00 | 207.33 | 1.24 | 1.00 |

| Contrast | Estimate | *SE* | *df* | *t* | $p_{adj}$ |
|---|---|---|---|---|---|
| 5–6-y CP – 8–11-y O | 0.00 | 0.00 | 174.60 | 0.30 | 1.00 |
| 5–6-y CP – 14–18-y O | 0.04 | 0.00 | 174.60 | 8.59 | **< .001** |
| 8–11-y CP – 14–18-y CP | 0.02 | 0.00 | 207.33 | 4.57 | **< .001** |
| 8–11-y CP – 5–6-y O | 0.01 | 0.00 | 174.60 | 1.68 | 1.00 |
| 8–11-y CP – 8–11-y O | 0.00 | 0.00 | 207.33 | 1.24 | 1.00 |
| 8–11-y CP – 14–18-y O | 0.04 | 0.00 | 207.33 | 12.48 | **< .001** |
| 14–18-y CP – 5–6-y O | -0.01 | 0.00 | 174.60 | -1.69 | 1.00 |
| 14–18-y CP – 8–11-y O | -0.01 | 0.00 | 207.33 | -3.33 | **.037** |
| 14–18-y CP – 14–18-y O | 0.03 | 0.00 | 207.33 | 7.91 | **< .001** |
| 5–6-y O – 8–11-y O | -0.00 | 0.00 | 174.60 | -0.77 | 1.00 |
| 5–6-y O – 14–18-y O | 0.04 | 0.00 | 174.60 | 7.52 | **< .001** |
| 8–11-y O – 14–18-y O | 0.04 | 0.00 | 207.33 | 11.25 | **< .001** |

*Note*. Degrees of freedom were determined using Satterthwaite's method. $P_{adj}$-values are Bonferroni corrected. *SE*= standard error, *df*= degrees of freedom. F = frontal, CP = centro-parietal, O = occipital.

Supplementary Table 24

*Type III analysis of variance table from a linear mixed-effects model on the effects of age group and slow oscillation topography on slow oscillation density*

| Predictor | Sum of squares | Mean square | $df_{Num}$ | $df_{Den}$ | *F* | *p* |
|---|---|---|---|---|---|---|
| Age group | 91.70 | 45.85 | 2 | 89.22 | 81.82 | < .001 |
| Topography | 3.21 | 1.61 | 2 | 206.49 | 2.87 | .059 |
| Age group*topography | 1.66 | 0.42 | 4 | 206.49 | 0.74 | .565 |

*Note*. $df_{Num}$ indicates the degrees of freedom numerator. $df_{Den}$ indicates the degrees of freedom denominator. Degrees of freedom were determined using Satterthwaite's method.

Supplementary Table 25

*Post-hoc comparisons for the effect of age group on slow oscillation density values based on estimated marginal means*

| Contrast | Estimate | *SE* | *df* | *t* | $p_{adj}$ |
|---|---|---|---|---|---|
| 5–6-year-olds – 8–11-year-olds | 0.89 | 0.27 | 61.98 | 3.25 | .006 |
| 5–6-year-olds – 14–18-year-olds | -0.47 | 0.27 | 61.98 | -1.72 | .271 |
| 8–11-year-olds – 14–18-year-olds | -1.36 | 0.11 | 206.49 | -12.77 | < .001 |

*Note*. Degrees of freedom were determined using Satterthwaite's method. $P_{adj}$-values are Bonferroni corrected. *SE*= standard error, *df*= degrees of freedom.

Supplementary Table 26

*Type III analysis of variance table from a linear mixed-effects model on the effects of age group and slow oscillation topography on slow oscillation amplitude*

| Predictor | Sum of squares | Mean square | $df_{Num}$ | $df_{Den}$ | $F$ | $p$ |
|---|---|---|---|---|---|---|
| Age group | 258,542.07 | 129,271.03 | 2 | 91.68 | 166.16 | < .001 |
| Topography | 174,735.63 | 87,367.81 | 2 | 208.07 | 112.30 | < .001 |
| Age group*topography | 100,654.90 | 25,163.72 | 4 | 208.07 | 32.34 | < .001 |

*Note.* $df_{Num}$ indicates the degrees of freedom numerator. $df_{Den}$ indicates the degrees of freedom denominator. Degrees of freedom were determined using Satterthwaite's method.

Supplementary Table 27

*Post-hoc comparisons for the slow oscillation amplitude interaction effect based on estimated marginal means*

| Contrast | Estimate | *SE* | *df* | *t* | $p_{adj}$ |
|---|---|---|---|---|---|
| 5–6-y F – 8–11-y F | -81.08 | 11.55 | 116.06 | -7.02 | **< .001** |
| 5–6-y F – 14–18-y F | -24.01 | 11.55 | 116.06 | -2.08 | 1.00 |
| 5–6-y F – 5–6-y CP | 1.44 | 8.05 | 208.07 | 0.18 | 1.00 |
| 5–6-y F – 8–11-y CP | -6.31 | 11.55 | 116.06 | -0.55 | 1.00 |
| 5–6-y F – 14–18-y CP | 60.76 | 11.55 | 116.06 | 5.26 | **< .001** |
| 5–6-y F – 5–6-y O | -7.38 | 8.05 | 208.07 | -0.92 | 1.00 |
| 5–6-y F – 8–11-y O | -11.98 | 11.55 | 116.06 | -1.04 | 1.00 |
| 5–6-y F – 14–18-y O | 80.61 | 11.55 | 116.06 | 6.98 | **< .001** |
| 8–11-y F – 14–18-y F | 57.06 | 6.87 | 208.07 | 8.31 | **< .001** |
| 8–11-y F – 5–6-y CP | 82.52 | 11.55 | 116.06 | 7.15 | **< .001** |
| 8–11-y F – 8–11-y CP | 74.76 | 6.87 | 208.07 | 10.89 | **< .001** |
| 8–11-y F – 14–18-y CP | 141.83 | 6.87 | 208.07 | 20.65 | **< .001** |
| 8–11-y F – 5–6-y O | 73.70 | 11.55 | 116.06 | 6.38 | **< .001** |
| 8–11-y F – 8–11-y O | 69.09 | 6.87 | 208.07 | 10.06 | **< .001** |
| 8–11-y F – 14–18-y O | 161.69 | 6.87 | 208.07 | 23.55 | **< .001** |
| 14–18-y F – 5–6-y CP | 25.46 | 11.55 | 116.06 | 2.20 | 1.00 |
| 14–18-y F – 8–11-y CP | 17.70 | 6.87 | 208.07 | 2.58 | .383 |
| 14–18-y F – 14–18-y CP | 84.77 | 6.87 | 208.07 | 12.35 | **< .001** |
| 14–18-y F – 5–6-y O | 16.64 | 11.55 | 116.06 | 1.44 | 1.00 |
| 14–18-y F – 8–11-y O | 12.03 | 6.87 | 208.07 | 1.75 | 1.00 |
| 14–18-y F – 14–18-y O | 104.63 | 6.87 | 208.07 | 15.24 | **< .001** |
| 5–6-y CP – 8–11-y CP | -7.76 | 11.55 | 116.06 | -0.67 | 1.00 |
| 5–6-y CP – 14–18-y CP | 59.31 | 11.55 | 116.06 | 5.14 | **< .001** |
| 5–6-y CP – 5–6-y O | -8.82 | 8.05 | 208.07 | -1.10 | 1.00 |

| Contrast | Estimate | *SE* | *df* | *t* | $p_{adj}$ |
|---|---|---|---|---|---|
| 5–6-y CP – 8–11-y O | -13.43 | 11.55 | 116.06 | -1.16 | 1.00 |
| 5–6-y CP – 14–18-y O | 79.17 | 11.55 | 116.06 | 6.86 | **< .001** |
| 8–11-y CP – 14–18-y CP | 67.07 | 6.87 | 208.07 | 9.77 | **< .001** |
| 8–11-y CP – 5–6-y O | -1.06 | 11.55 | 116.06 | -0.09 | 1.00 |
| 8–11-y CP – 8–11-y O | -5.67 | 6.87 | 208.07 | -0.83 | 1.00 |
| 8–11-y CP – 14–18-y O | 86.93 | 6.87 | 208.07 | 12.66 | **< .001** |
| 14–18-y CP – 5–6-y O | -68.13 | 11.55 | 116.06 | -5.90 | **< .001** |
| 14–18-y CP – 8–11-y O | -72.74 | 6.87 | 208.07 | - | **< .001** |
| 14–18-y CP – 14–18-y O | 19.86 | 6.87 | 208.07 | 2.89 | .152 |
| 5–6-y O – 8–11-y O | -4.60 | 11.55 | 116.06 | -0.40 | 1.00 |
| 5–6-y O – 14–18-y O | 87.99 | 11.55 | 116.06 | 7.62 | **< .001** |
| 8–11-y O – 14–18-y O | 92.60 | 6.87 | 208.07 | 13.48 | **< .001** |

*Note*. Degrees of freedom were determined using Satterthwaite's method. $P_{adj}$-values are Bonferroni corrected. *SE*= standard error, *df*= degrees of freedom. F = frontal, CP = centro-parietal, O = occipital.

Supplementary Table 28

*Results of the generalized linear mixed-effects model on the association between the fast sleep spindle maturity component and frontal slow oscillation-development-specific fast spindle modulation strength*

| Predictor | Estimate | Standard error | *t* | *p* |
|---|---|---|---|---|
| (Intercept) | -1.85 | 0.07 | -25.13 | < .001 |
| Fast sleep spindle maturity component | 0.36 | 0.07 | 5.40 | < .001 |
| Slow oscillation maturity score | 0.09 | 0.07 | 1.29 | .197 |

*Note*. Model was fit with gamma errors and a log link function.

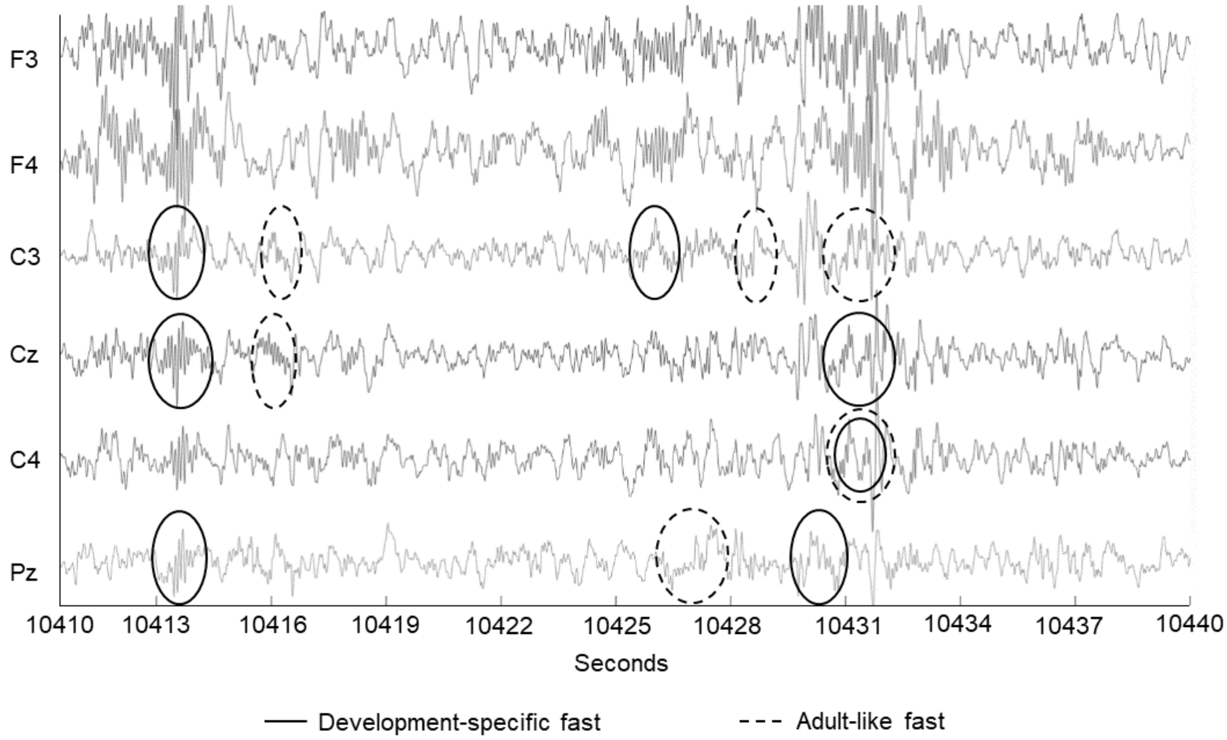


*Supplementary Figure 1.* Example 30-second raw EEG window from a 6-year-old child with events that were detected as development-specific fast (solid circles) and adult-like fast (dashed circles) sleep spindles.

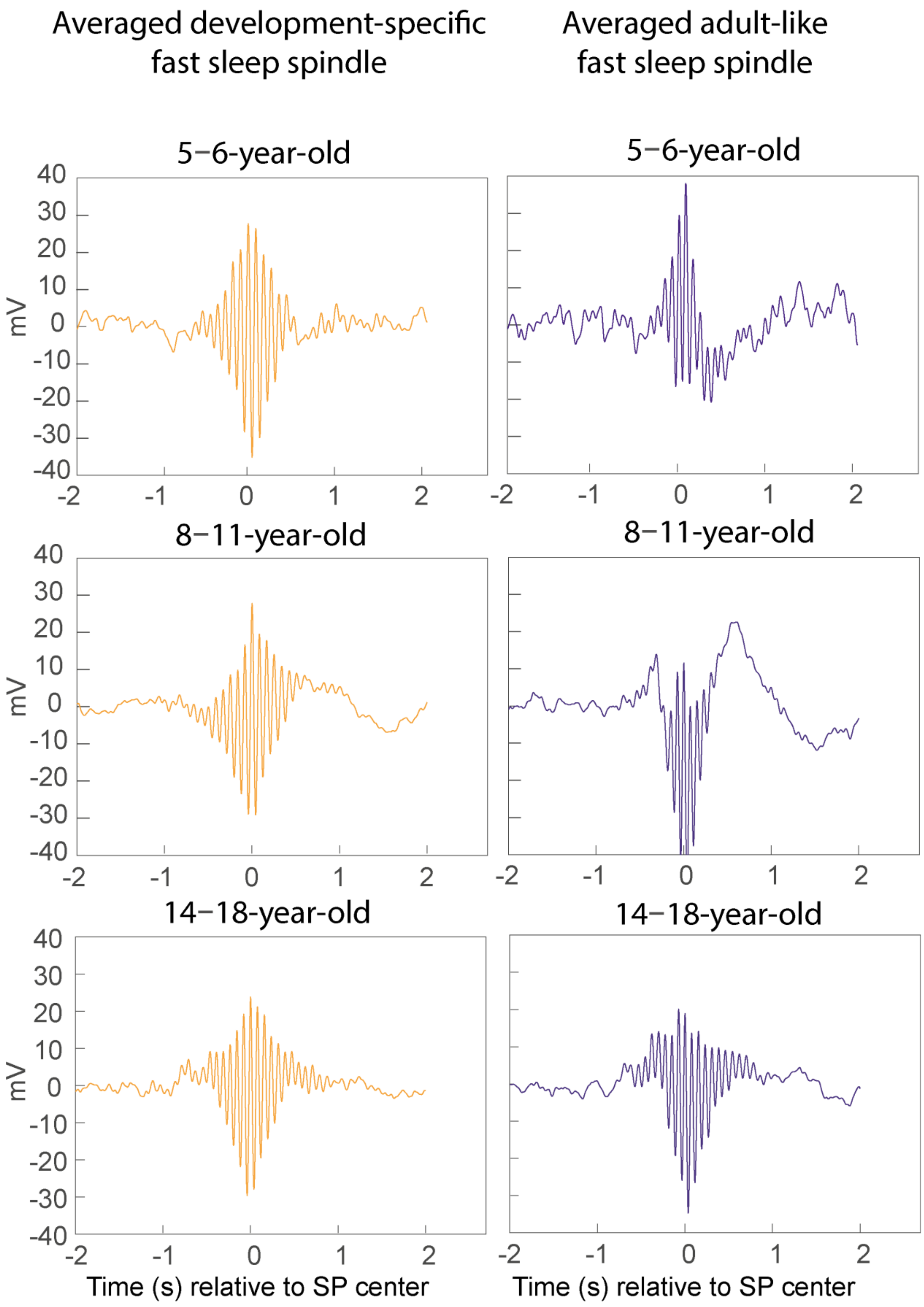


*Supplementary Figure 2.* Appearance of averaged detected development-specific and adult-like fast sleep spindles for one 6-year-old participant and one participant who underwent EEG at 9 years of age and again at 15 years of age.

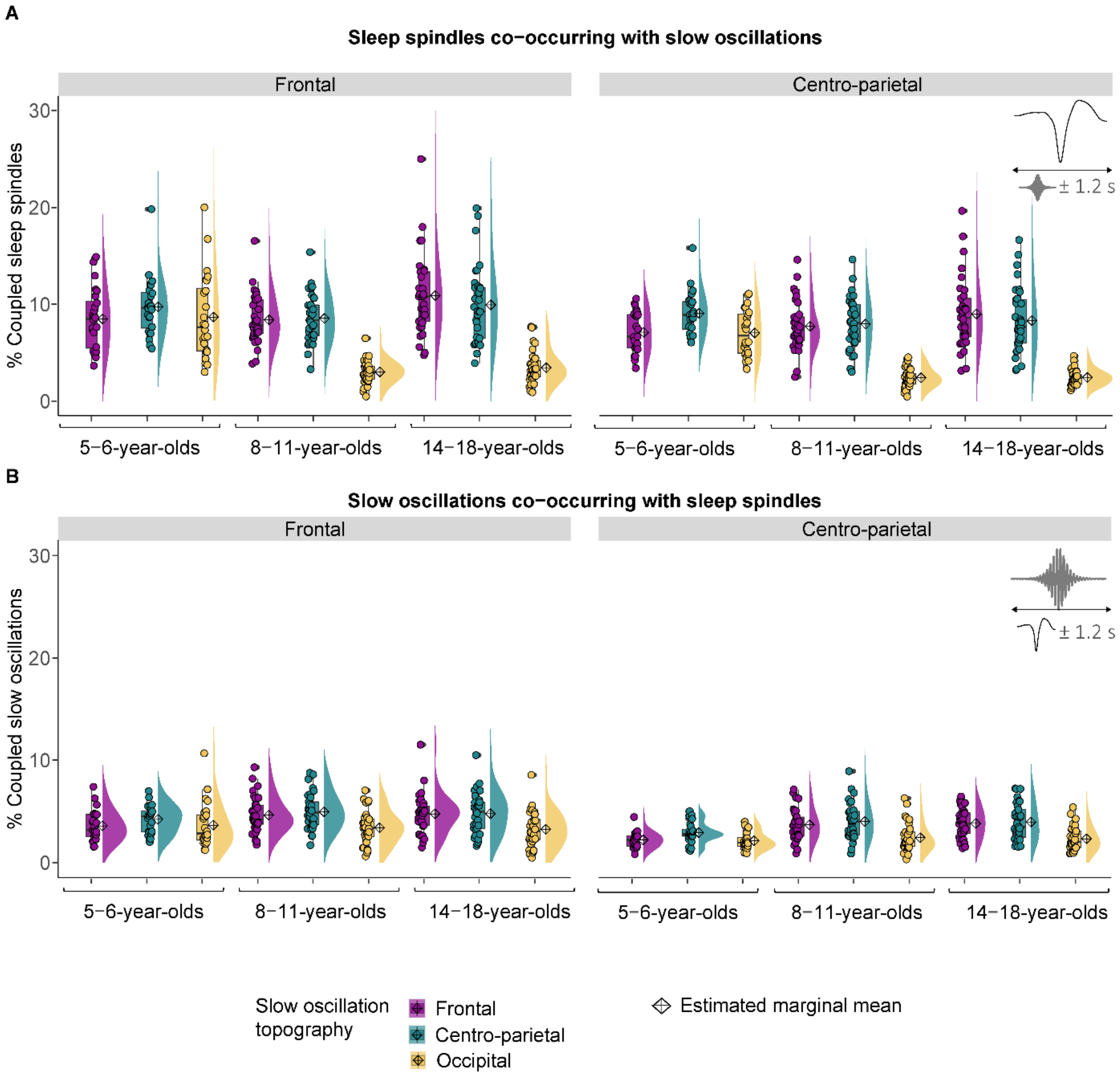


*Supplementary Figure 3.* Co-occurrence data of slow frontal and development-specific fast centro-parietal sleep spindles and slow oscillations in different topographical locations. (A) Percentage of spindle centers occurring within ± 1.2 s around the slow oscillation down peak. (B) Percentage of slow oscillation down peaks occurring within ± 1.2 s around the spindle centers. Diamonds represent estimated marginal means from linear mixed-effects models depicted in Supplementary Tables 2-5.

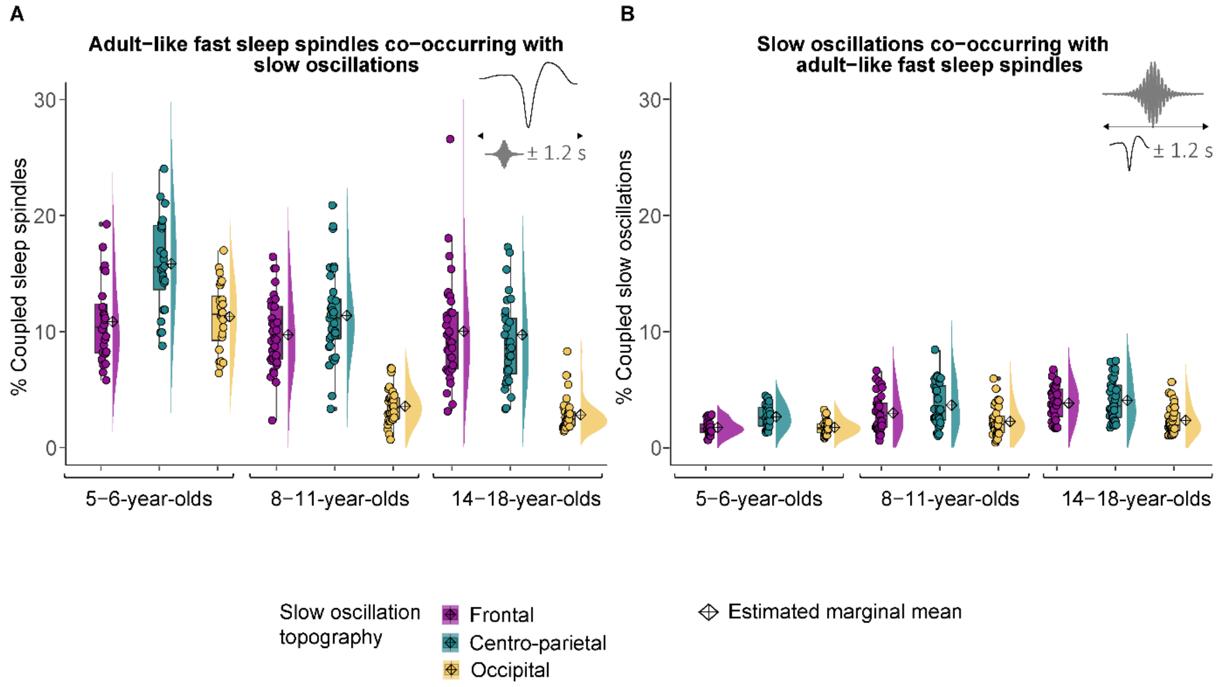


*Supplementary Figure 4.* Co-occurrence data of adult-like fast sleep spindles and slow oscillations in different topographical locations. (A) Percentage of spindle centers occurring within ± 1.2 s around the slow oscillation down peak. (B) Percentage of slow oscillation down peaks occurring within ± 1.2 s around the spindle centers. Diamonds represent estimated marginal means from linear mixed-effects models depicted in Supplementary Tables 6 and 7.

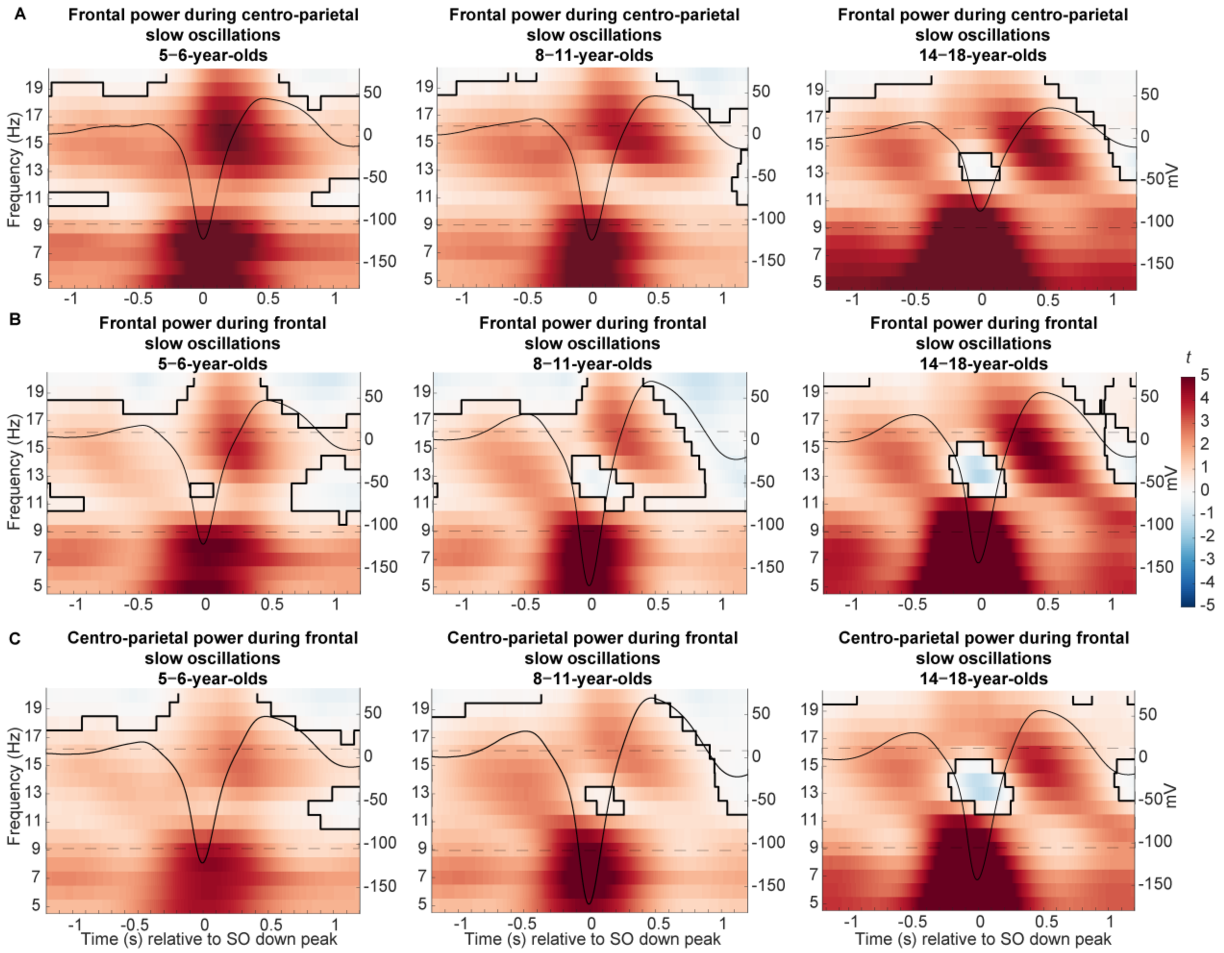


*Supplementary Figure 5.* (A-B) Frontal and (C) centro-parietal power differences between trials with and without (A) centro-parietal and (B-C) frontal slow oscillations (in *t*-score units). Significant clusters are outlined in black (cluster-based permutation test, cluster $\alpha < .05$, two-sided test). The average slow oscillation for each age group is plotted onto the power differences in black to illustrate the relation to slow oscillation phase (scale in mV on the right y-axis of each plot). The sleep spindle frequency range is highlighted by the dashed window.

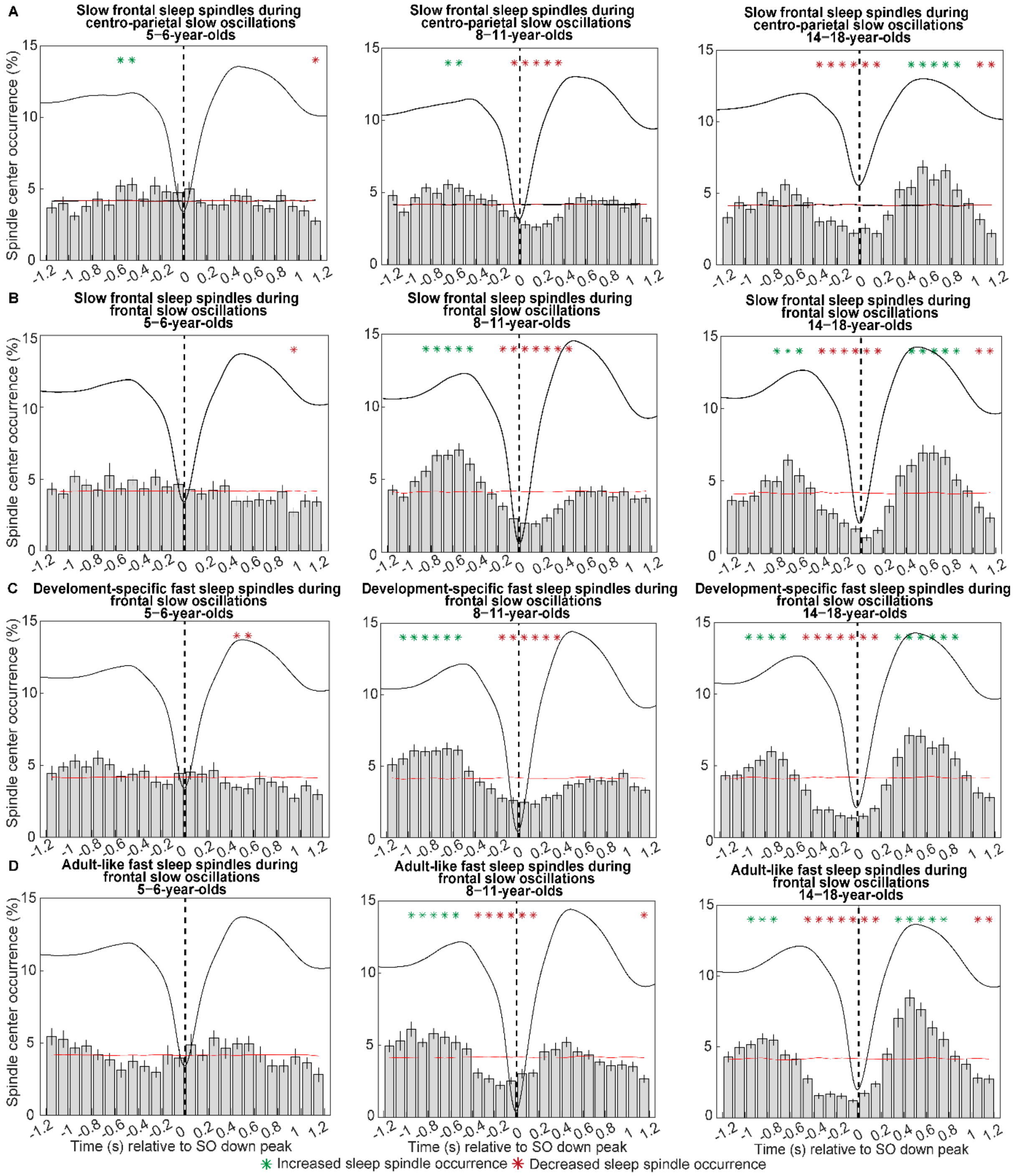


*Supplementary Figure 6.* Peri-event time histograms for (A-B) slow frontal (C) development-specific fast centro-parietal, and (D) adult-like fast spindles showing the proportion of sleep spindles occurring within 100 ms bins during (A) centro-parietal (B-D) frontal slow oscillations. Error bars represent standard errors. Green asterisks mark increased spindle occurrence (positive cluster, cluster α < .05, two-sided test) and red asterisks mark decreased spindle occurrence (negative cluster, cluster α < .05, two-sided test) compared to random occurrence (horizontal line). The dashed vertical line indicates the slow oscillation down peak. The average slow oscillation of each age group is shown in black to illustrate the relation to the slow oscillation phase.

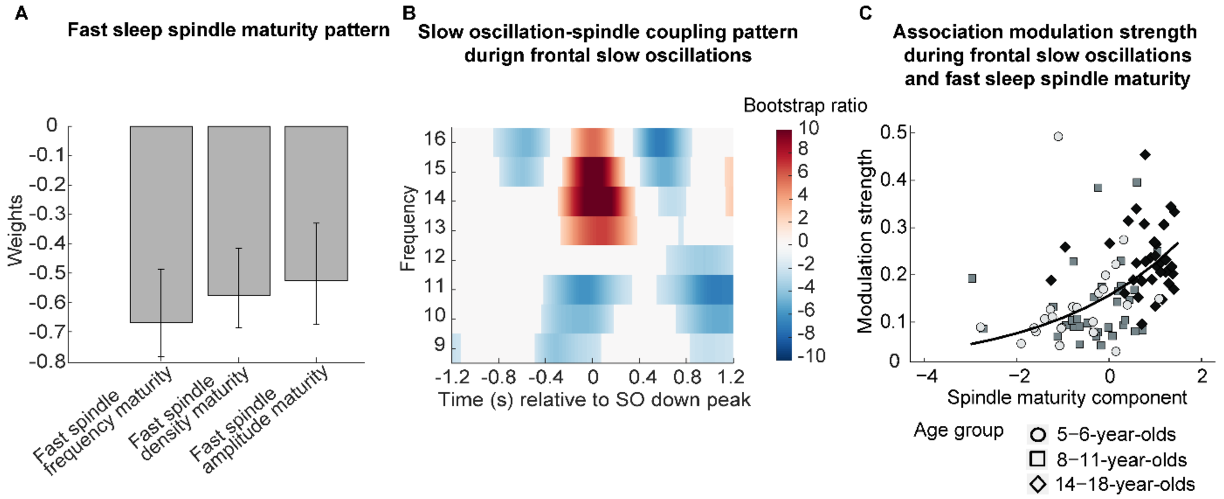


*Supplementary Figure 7.* Association between fast spindle maturity and frontal slow oscillation-development-specific fast spindle coupling. (A-B) Results from a partial least squares correlation revealed one (A) fast spindle maturity profile significantly associated with (B) the frontal slow oscillation-spindle coupling pattern. (A) Weights of the first singular vector dimension of the fast spindle maturity scores. Error bars represent 95% bootstrap confidence intervals. (B) Weights of the first singular vector dimension of the slow oscillation-spindle coupling pattern by means of bootstrap ratios. Only values >1.96 and < -1.96 are colored. (C) Scatterplot of the association between the modulation strength (KL divergence) of development-specific fast centro-parietal SPs during frontal SOs and the fast spindle maturity component. The curved line represents the prediction from the generalized linear mixed-effects model for the simple effect of the spindle maturity component (Supplementary Table 28). For visualization purposes, the three age groups are indicated by different shapes and colors.
